# Diel Oscillations of Particulate Metabolites Reflect Synchronized Microbial Activity in the North Pacific Subtropical Gyre

**DOI:** 10.1101/2020.05.09.086173

**Authors:** Angela K. Boysen, Laura T. Carlson, Bryndan P. Durham, Ryan D. Groussman, Frank O. Aylward, François Ribalet, Katherine R. Heal, Edward F. DeLong, E. Virginia Armbrust, Anitra E. Ingalls

## Abstract

Light is the primary input of energy into the sunlit ocean, driving daily oscillations in metabolism of primary producers. The consequences of this solar forcing have implications for the whole microbial community, yet *in situ* measurements of metabolites, direct products of cellular activity, over the diel cycle are scarce. We evaluated community-level biochemical consequences of diel oscillations in the North Pacific Subtropical Gyre by quantifying 79 metabolites in particulate organic matter in surface waters every four hours over eight days. Total particulate metabolite concentration peaked at dusk, even when normalized to biomass estimates. The concentrations of 70% of individual metabolites exhibited 24-hour periodicity. Despite the diverse organisms that use them, primary metabolites involved in anabolic processes and redox maintenance had significant 24-hour periodicity. Osmolytes exhibited the largest diel oscillations, implying rapid turnover and metabolic roles beyond cell turgor maintenance. Metatranscriptome analysis revealed the taxa involved in production and consumption of some metabolites, including the osmolyte trehalose. This compound displayed the largest diel oscillations in abundance and was likely produced by the nitrogen-fixing cyanobacterium *Crocosphaera* for energy storage. These findings demonstrate that paired measurements of particulate metabolites and transcripts resolve strategies microbes use to manage daily energy and redox oscillations.

## Introduction

Light is a powerful forcing on metabolism in the surface ocean, acting at the molecular level to drive global biogeochemical cycles(1). In surface waters near Station ALOHA (22º 45’N, 158º W) in the North Pacific Subtropical Gyre (NPSG), the marine plankton community responds to diel forcing, either directly or indirectly, as demonstrated by daily oscillations in particulate organic carbon (POC)(2), cell division(3), gross primary production, net community production(4), grazing(5), viral infection(6), and nitrogen fixation(7). Genes associated with a wide variety of cellular processes also exhibit diel oscillations in transcript abundance, reflecting the capture of light energy and its conversion to chemical energy during daylight, a process that fuels metabolism over a 24 hour period(8–12). Temporal partitioning of anabolism and catabolism creates diel patterns in the macromolecular composition of the community(12–15).

Though most POC in the surface ocean is made up of macromolecules(16,17), the suite of small molecules (< 800 daltons) produced within cells helps shape the internal and external chemical environment of the plankton community. Small molecules are likely responsible for setting dependencies among different taxa, yet an inventory of these compounds and the plasticity of their concentrations remain largely unknown(18). Comprehensive measurements of intracellular small biomolecules, or metabolites, present in marine microbial communities are scarce and the suite of compounds detected is strongly biased by the methods employed(19). Small polar molecules, in particular, are rarely measured although they are the main component of the aqueous cytosol. Intracellular metabolite profiles of model marine microbes are taxon-specific and respond to environmental perturbations, including diel oscillations in available light(20). Many of these metabolites have not yet been measured in plankton communities and some are without annotated biosynthetic or catabolic pathways(21–27). Thus, a comprehensive inventory of intracellular metabolites will facilitate a deeper understanding of marine microbial physiology and interactions that drive ecosystem diversity and activity(28,29).

Here we measured particulate metabolite concentrations in samples collected from surface waters near Station ALOHA during eight diel cycles. These data provide an inventory of metabolites in the oligotrophic surface ocean and show the metabolic consequences of the diel cycle. We paired observations of metabolites with gene expression data, POC, and flow cytometry (FCM) measurements. We inferred community physiology over the diel cycle to predict how environmental conditions produce a particular chemical environment within natural populations of marine plankton.

We find that the molar concentration of 70% of our targeted metabolites oscillate with 24-hour periodicity, reflecting large scale community synchrony. Our analysis identifies diel oscillations in compounds that play important roles in managing light-induced redox reactions and biosynthesis of building blocks and energy stores. These compounds are ultimately conduits of energy and nutrients through the microbial ecosystem as they are exchanged between diverse organisms, with repercussions for community diversity and function(30–32). By pairing metabolite data with metatranscriptomes, we identify potential metabolic strategies that organisms deploy for coping with redox oscillations induced by an oscillating energy supply.

## Materials and Methods

### Sample Collection

Samples were collected on the R/V Kilo Moana in the NPSG (near 24.5º N, 156.5º W) every four hours for two sampling periods in summer 2015 (period one: July 26, 6:00 – July 30, 6:00; period two: July 31 18:00 – August 3, 18:00). To limit variability unrelated to solar forcing, we conducted Lagrangian sampling following two drifters in an anticyclonic eddy(7). Samples for particulate metabolites and transcripts were collected from 15 m water depth using Niskin bottles attached to a conductivity, temperature, depth array (CTD). Ancillary measurements for nutrients and heterotrophic bacterial abundance (reported in Wilson et. al. 2017) were collected and analyzed with standard Hawaii Ocean Time-series protocols (http://hahana.soest.hawaii.edu/index.html).

### Bulk and taxa-specific carbon biomass

POC concentrations were derived from particulate beam attenuation at 660 nm, as in White et al.(2). Particle absorption was calibrated against discrete POC samples taken near dawn and dusk. Discrete POC and particulate nitrogen (PN) samples were collected by filtration of the ship’s underway flow through seawater onto combusted GFF filters. Analysis is described in the supplemental methods.

Continuous underway flow cytometry (SeaFlow)(33) was used to count *Prochlorococcus, Synechococcus*, picoeukaryotes (eukaryotic phytoplankton 2–4 µm in size), and *Crocosphaera*. These data were supplemented with discrete flow cytometry sample analysis as in Wilson et al.(7). Cell diameters of individual cells were estimated from light scatter by the application of Mie theory to a simplified optical model and converted to carbon quotas assuming spherical particles, as described in Ribalet et al.(34). Carbon biomass was estimated by multiplying cell abundance by carbon quotas.

### Metabolite extraction, data acquisition, and processing

Metabolite samples were collected in triplicate at each time point by filtering 3.5 L of seawater onto a 47 mm 0.2 *µ*m PTFE (Omnipore) filter using a peristaltic pump, polycarbonate filter holder, and Masterflex PharMed BPT tubing (Cole-Parmer). Filters were frozen in liquid nitrogen immediately after filtration and stored at -80 °C. Metabolite extractions employed a modified Bligh-Dyer method(19,27,35), resulting in aqueous and organic soluble metabolites with isotope-labeled extraction and injection internal standards added to both fractions (Table S1, Supplemental Methods). Unused filters served as methodological extraction blanks.

Metabolomics data were collected by paired liquid-chromatography mass-spectrometry (LC-MS) using both hydrophilic liquid interaction chromatography and reversed phase chromatography with a Waters Acquity I-Class UPLC and a Waters Xevo TQ-S triple quadrupole with electrospray ionization in selected reaction monitoring mode with polarity switching, targeting over 200 compounds(19). The software Skyline was used to integrate LC-MS peaks(36) and data were normalized using best-matched internal standard normalization(19). A subset of this data are presented in Durham et al. (2019) and Muratore et al. (2020)(14,27).

Metabolites with isotopologue internal standards were quantified in all samples (Table S1). Trehalose, sucrose, and 2,3-dihydroxypropane-1-sulfonate (DHPS) were quantified with standard additions. For all other metabolites, concentration (pmol L^-1^) was calculated from injections of known concentrations of authentic standards in both water and a representative matrix to correct for ion suppression. Dimethylsulfoniopropionate (DMSP) loss is known to occur during methanol-based extractions so concentrations are considered a minimum estimate(37). Details are in the supplemental methods.

### Metatranscriptome data acquisition and processing

Whole community transcript data are referred to here as prokaryotic transcript data, as they were enriched in bacterial and archaeal RNA. These metatranscriptome samples were collected on 0.2 µm filters simultaneously with the metabolomic data reported here, as previously reported in Wilson et al.(7) and Aylward et al.(6). The metatranscriptome sequence reads were quality trimmed, end-joined, mapped, and quantified with molecular standards.

Metatranscriptome sequence reads were mapped to the ALOHA gene catalog(38) using LAST v 959(39), and transcript count normalization, leveraging the molecular standards described in Gifford et al.(40). Sequence reads were summed if assigned to the same taxonomic order and Kyoto Encyclopedia of Genes and Genomes (KEGG) orthologue(41).

Poly-A+ selected transcript data (referred to here as eukaryotic transcript data) are from the metatranscriptomes presented in Durham et al.(27). These samples were collected on 0.2 µm filters concurrently with the metabolomic samples and include only the first sampling period. Quality-controlled short reads were assembled using Trinity *de novo* transcriptome assembler version 2.3.2(42). Using DIAMOND v 0.9.18(43), assembled contigs were aligned to a reference sequence database of marine organisms (MarineRefII reference database, http://roseobase.org/data/, with additions listed in Table S2). Taxonomy was assigned with DIAMOND by using the top 10% of hits with e-value scores below 10^−5^ to estimate the Lowest Common Ancestor of each contig. We assigned putative function using hmmsearch (from HMMER 3.1b2(44), minimum bitscore 30) to find the best-scoring KEGG gene family from KOfam (ver. 2019-03-20) (45). Contig abundances were quantified by mapping the paired reads to the assemblies with kallisto(46). Sequence reads assigned to the same taxonomic group and KEGG ortholog were summed and normalized to the total read pool of the taxonomic group. Details are provided in the supplemental methods.

Metabolites and transcripts were associated with one another using the KEGG database as a scaffold to match metabolites with transcripts coding for enzymes that directly use or produce those metabolites. The R package KEGGREST(47) was used to access the KEGG database followed by manual curation of these matches.

### Detecting Periodicity

Diel periodicity was evaluated for all signals using Rhythmicity Analysis Incorporating Non-parametric Methods (RAIN)(6,7,48). Metabolites and transcripts were considered significantly periodic if they had a false discovery rate (fdr)(49) corrected *p*-value < 0.05. For each significantly oscillating signal, the time of peak abundance was estimated by fitting a periodic function (supplemental methods), though we recognize the precision of these peak times is limited by our sampling resolution. Diel periodicity in metabolites was identified for the two different sampling periods independently and jointly.

### Phytoplankton culture conditions

Cultures of phytoplankton were grown in combusted borosilicate tubes in diurnal incubators with a 12:12 light:dark cycle. Samples for metabolomics were collected by gentle filtration onto 0.2 µm Durapore filters using combusted borosilicate filter towers. *Crocosphaera watsonii* strain WH8501 was grown at 27 °C with 50 µmol photons m^-2^ s^-1^ in YBC-II artificial seawater medium(50) supplemented with 0.9 mM nitrate; cells were collected just before the lights turned on and just after the lights turned off during exponential phase. Cells were enumerated via a Beckman Z2 Coulter Counter. *Prochlorococcus* MIT1314 (HLII clade(51)) were grown at 20 °C with 20 µmol photons m^-2^ s^-1^ in Pro99 media(52) prepared with Turks Island Salt Solution and supplemented with 6 mM sterile sodium bicarbonate and 1 mM N-Tris(hydroxymethyl)methyl-3-aminopropanesulfonic acid(53). *Prochlorococcus* cells were collected 6 hours into the light period during exponential phase and enumerated using the flow cytometer BD Influx cell sorter. Axenicity of *Prochlorococcus* cultures was verified regularly with SYBR-staining and FCM and plating on bacterial ½ YTSS agar.

## Results

### Oscillatory dynamics of the phytoplankton community

Our sampling targeted an anticyclonic eddy to facilitate Lagrangian sampling, and was characterized by warm, nutrient-deplete surface waters typical of the persistently oligotrophic NPSG(5,54) (Table 1). Photosynthetic picoeukaryotes, *Prochlorococcus*, and *Crocosphaera* contributed substantially to phytoplankton biomass(7) (Figure 1). POC, which includes bulk community biomass, and phytoplankton-specific biomass oscillated with significant 24-hour periodicity (Figure 1). Cell abundances and total biomass of *Prochlorococcus* and *Crocosphaera* populations increased between the first and second sampling periods (Table 1). Wind speed also increased between the first and second sampling periods, resulting in an increase in the mixed layer depth from 21 ± 5 to 36 ± 6 m. Additionally, we observed a decrease in the number of significantly diel metabolite oscillations during the second sampling interval, from 55 to 9 (Table S3). This change was likely related to the deepening of the mixed layer; however, we have insufficient evidence to investigate this hypothesis further. We therefore focus on data collected during the first sampling period.

**Table 1.**
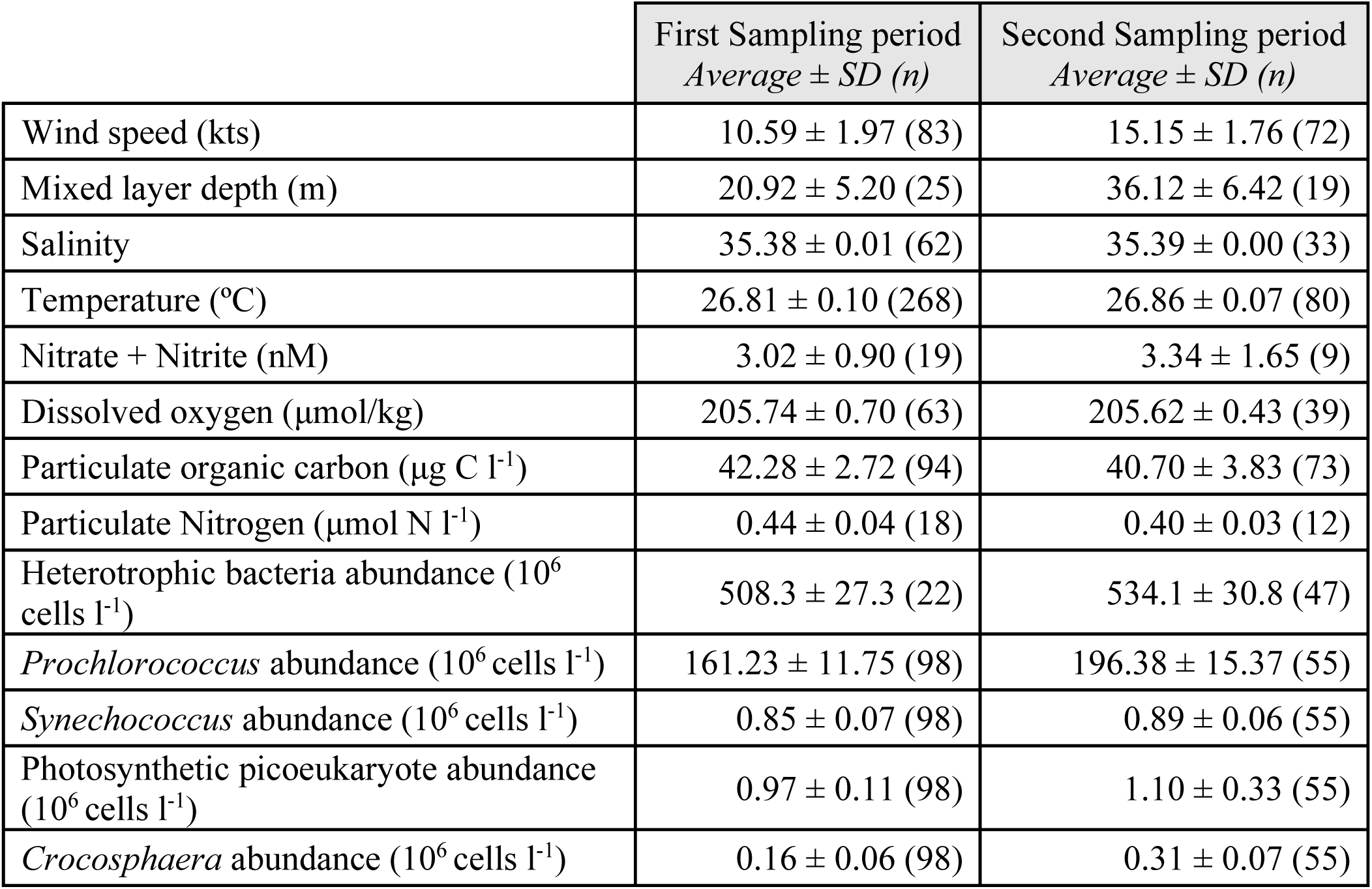
Wind speed and surface mixed layer physical and biological variables over the two sampling periods. Salinity, temperature, and dissolved oxygen (corrected with bottle measurements) are from the CTD between 13-17 m. N+N and heterotrophic bacteria abundance are measured from discrete samples at 15 m. Particulate organic carbon (from underway beam attenuation), particulate nitrogen, *Prochlorococcus, Synechococcus*, photosynthetic picoeukaryotes, and *Crocosphaera* are measured from the ship-underway water intake near 7 m.

|  | First Sampling period<br>*Average ± SD (n)* | Second Sampling period<br>*Average ± SD (n)* |
| --- | --- | --- |
| Wind speed (kts) | 10.59 ± 1.97 (83) | 15.15 ± 1.76 (72) |
| Mixed layer depth (m) | 20.92 ± 5.20 (25) | 36.12 ± 6.42 (19) |
| Salinity | 35.38 ± 0.01 (62) | 35.39 ± 0.00 (33) |
| Temperature (°C) | 26.81 ± 0.10 (268) | 26.86 ± 0.07 (80) |
| Nitrate + Nitrite (nM) | 3.02 ± 0.90 (19) | 3.34 ± 1.65 (9) |
| Dissolved oxygen (μmol/kg) | 205.74 ± 0.70 (63) | 205.62 ± 0.43 (39) |
| Particulate organic carbon (μg C l<sup>-1</sup>) | 42.28 ± 2.72 (94) | 40.70 ± 3.83 (73) |
| Particulate Nitrogen (μmol N l<sup>-1</sup>) | 0.44 ± 0.04 (18) | 0.40 ± 0.03 (12) |
| Heterotrophic bacteria abundance (10<sup>6</sup> cells l<sup>-1</sup>) | 508.3 ± 27.3 (22) | 534.1 ± 30.8 (47) |
| *Prochlorococcus* abundance (10<sup>6</sup> cells l<sup>-1</sup>) | 161.23 ± 11.75 (98) | 196.38 ± 15.37 (55) |
| *Synechococcus* abundance (10<sup>6</sup> cells l<sup>-1</sup>) | 0.85 ± 0.07 (98) | 0.89 ± 0.06 (55) |
| Photosynthetic picoeukaryote abundance (10<sup>6</sup> cells l<sup>-1</sup>) | 0.97 ± 0.11 (98) | 1.10 ± 0.33 (55) |
| *Crocospaera* abundance (10<sup>6</sup> cells l<sup>-1</sup>) | 0.16 ± 0.06 (98) | 0.31 ± 0.07 (55) |

**Figure 1.**
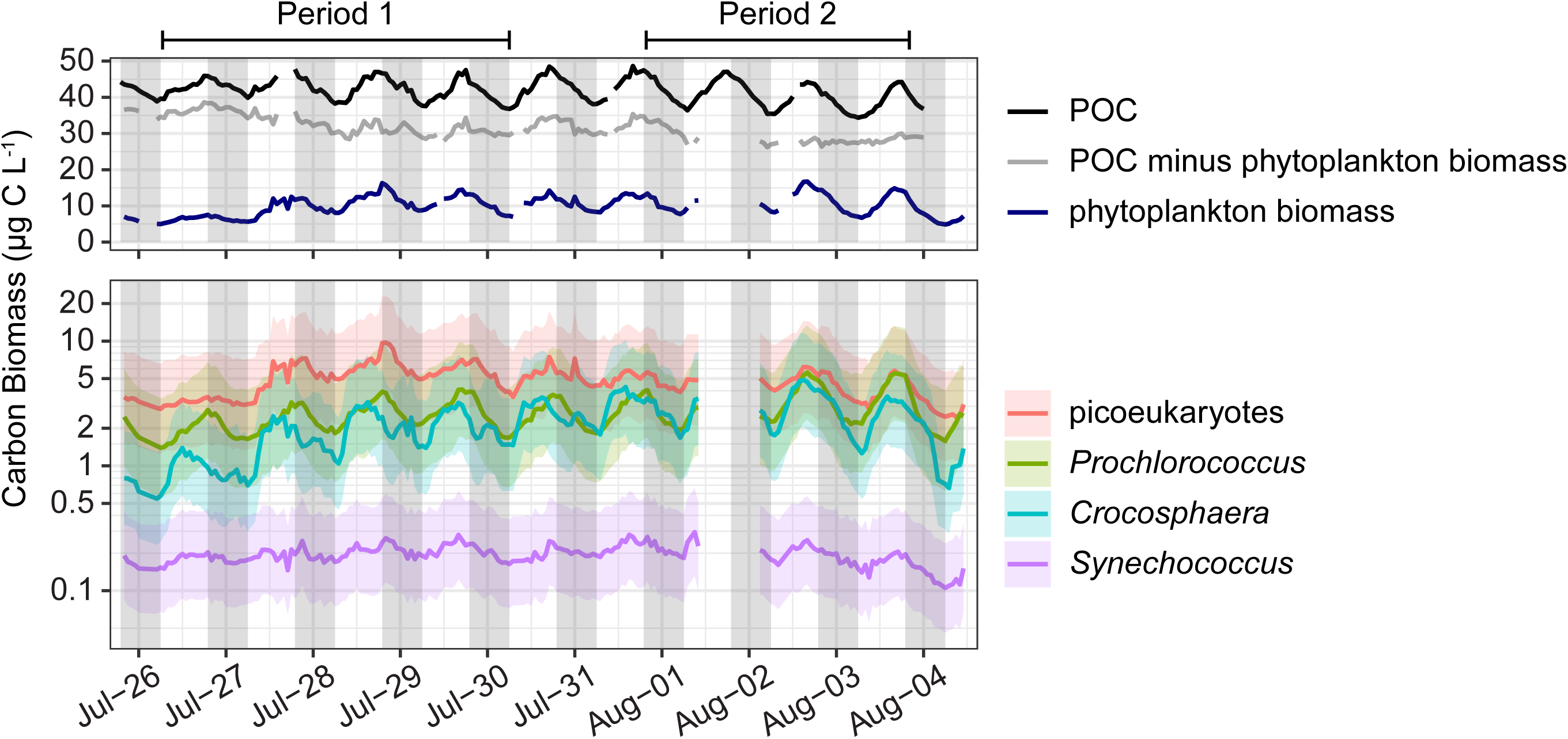
Top: Hourly averages of POC from beam attenuation (black line), total phytoplankton carbon biomass from flow cytometry (phytoplankton biomass, navy line), and the difference between the two (grey line). Bottom: Hourly averages of population specific carbon biomass of *Prochlorococcus, Synechococcus, Crocosphaera*, and photosynthetic picoeukaryotes (defined here as 2-4 µm) from flow cytometry, with shaded area representing the 95% confidence interval. Breaks in the lines are due to short periods of instrument malfunction. The two sampling periods referred to in the text are indicated above the figure.

### Metabolite Inventory

A total of 79 targeted metabolites were detected across samples (Table S3). Total particulate metabolite concentration increased during the day and decreased at night, regardless of whether normalized to POC or PN (Figure 2). The most abundant compounds were osmolytes, like glycine betaine (GBT), homarine, DHPS, and DMSP; nucleobases (particularly guanine); and amino acids related to nitrogen metabolism, such as glutamic acid and glutamine (Table S3, Figure 2). At dusk, quantified metabolites totaled 1.7 ± 0.2% of POC and 3.1 ± 0.6% of particulate nitrogen (Figure 2), with free nucleobases and amino acids representing substantial pools of total cellular nitrogen (Table S4).

**Figure 2.**
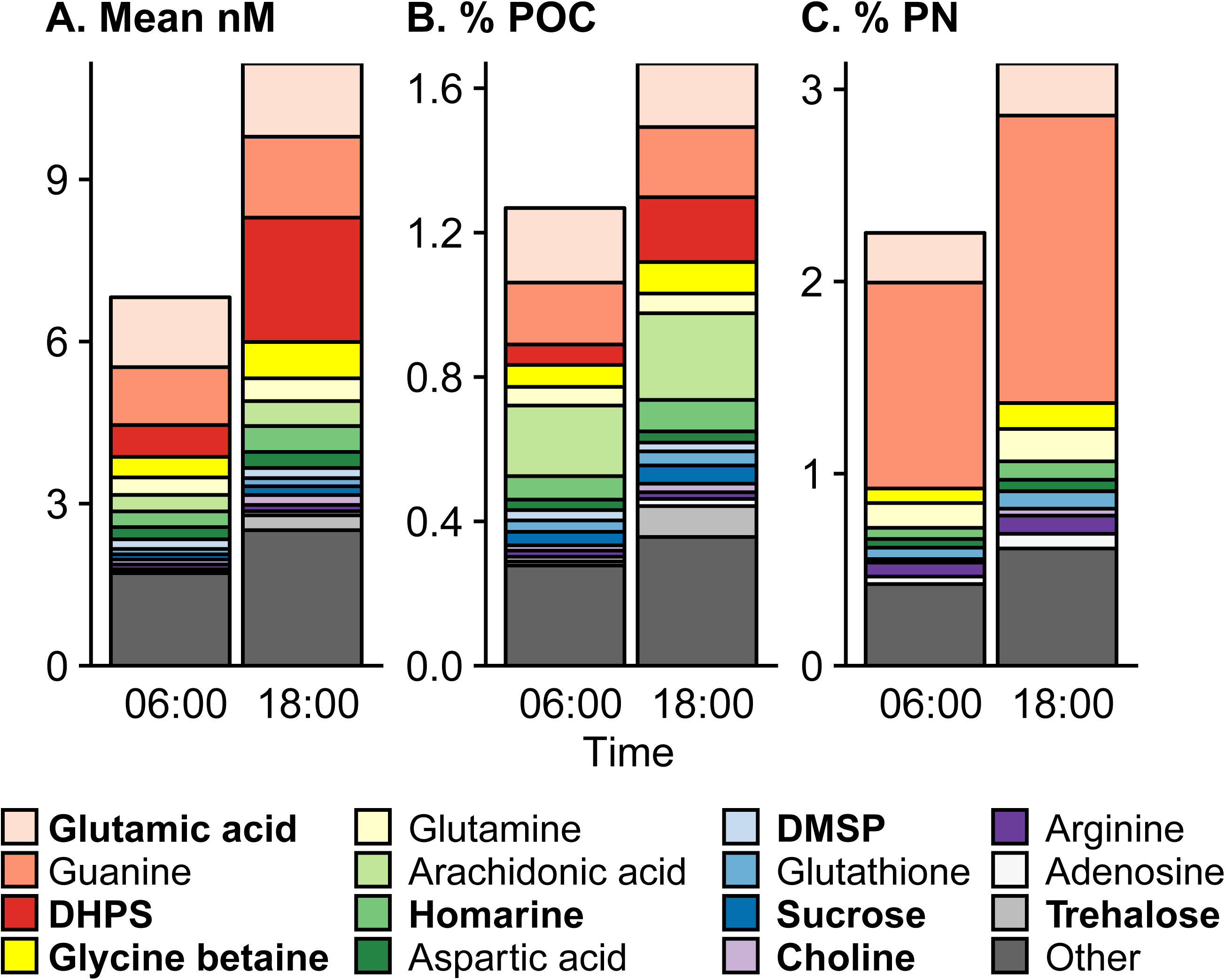
Average targeted metabolite composition at dawn (06:00) and dusk (18:00) from July 26th – July 28th (n = 9 for each time point), shown as the estimated particulate metabolite concentration (A), the percent of particulate organic carbon (B), and the percent of the particulate nitrogen (C). “Other” contains the sum of the rest of the metabolites (64 compounds). Osmolytes are in bold. Note the different y-axis scales.

Multivariate analyses were used to determine if time of day influenced the community metabolome. NMDS analysis shows that samples collected at different times were significantly different. Samples collected near sunrise (6:00) were more similar to one another than those collected at other times of day and are most dissimilar to samples collected near sunset (ANOSIM, *R* = 0.19, *p* = 0.001, Figure S1, Table S5).

### Metabolite diel periodicity

To determine whether metabolite oscillations were driven by changes in biomass or by changing cell physiology resulting in changes in the intracellular concentration, we calculated concentrations relative to water volume filtered, resulting in values proportional to molar concentration (nmol L^-1^), and to POC, resulting in values proportional to nmol per *µ*mol POC. Bulk and individual metabolite concentrations oscillated with respect to both normalizations (Figure 2, Figure 3A). The molar concentration (nmol L^-1^) of 55 metabolites (70%) had significant 24-hour oscillations, with 26 reaching a maxima in concentration within two hours of 18:00 and 20 reaching their peak concentration within 2 hours of 14:00 (Figure 3, Table S3). When normalized to POC (nmol *µ*mol POC^-1^), 37 compounds (47%) showed diel oscillations (Table S3), and the mean time of peak concentration shifted to earlier in the afternoon (Figure 3A). POC reflects total community biomass and detritus, so to avoid assumptions of metabolite source, we present molar concentrations throughout except where metabolite source can be constrained to a specific phytoplankton, in which case we present metabolite concentration normalized to the cell number or biomass of the source organism.

**Figure 3.**
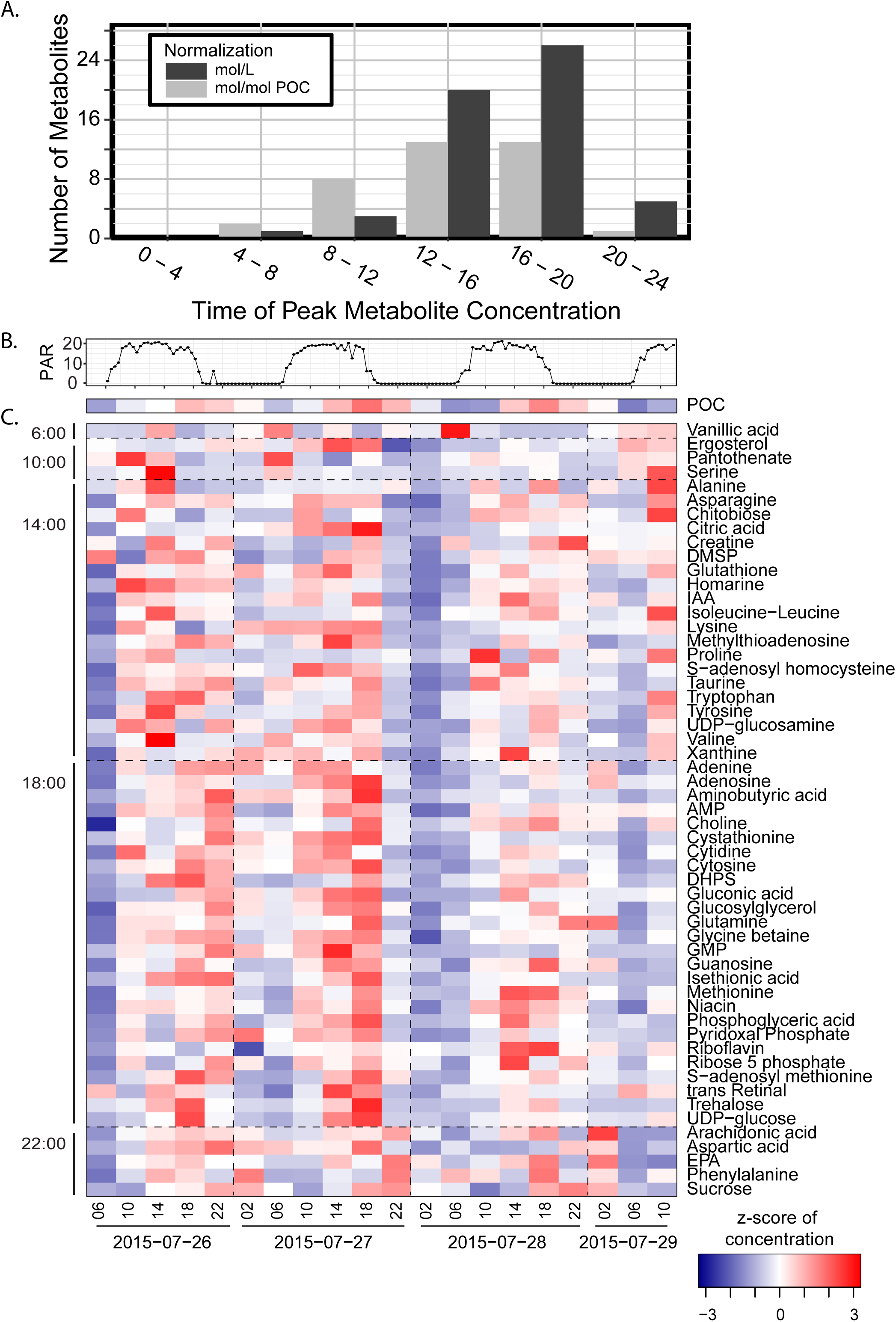
Time of day that significantly diel compounds peak in the first sampling period (A). Surface light (photosynthetically active radiation, PAR, x10 nmol photon m-2 s-1) (B). Heat map showing the z-score standardized concentrations of POC and of metabolites (nmol L-1) determined to be significantly diel in the first sampling period, arranged by time of peak concentration (C).

Metabolites with significant oscillations had daily fold changes ranging from 2 to 12.8, all of which exceeded the 1.2- and 1.8-fold changes of POC and the sum of FCM phytoplankton biomass, respectively (Figure 4). The disaccharides trehalose and sucrose displayed the most robust oscillations (*p*-value <1×10^−13^, Figures 4, 5). Trehalose and sucrose are known osmolytes, and nearly all other identified osmolytes (9/10) showed diel oscillations (Figures 4, 6, Table S3). Glutamic acid is the only known osmolyte that did not have a significant oscillation in molar concentration (Table S3).

**Figure 4.**
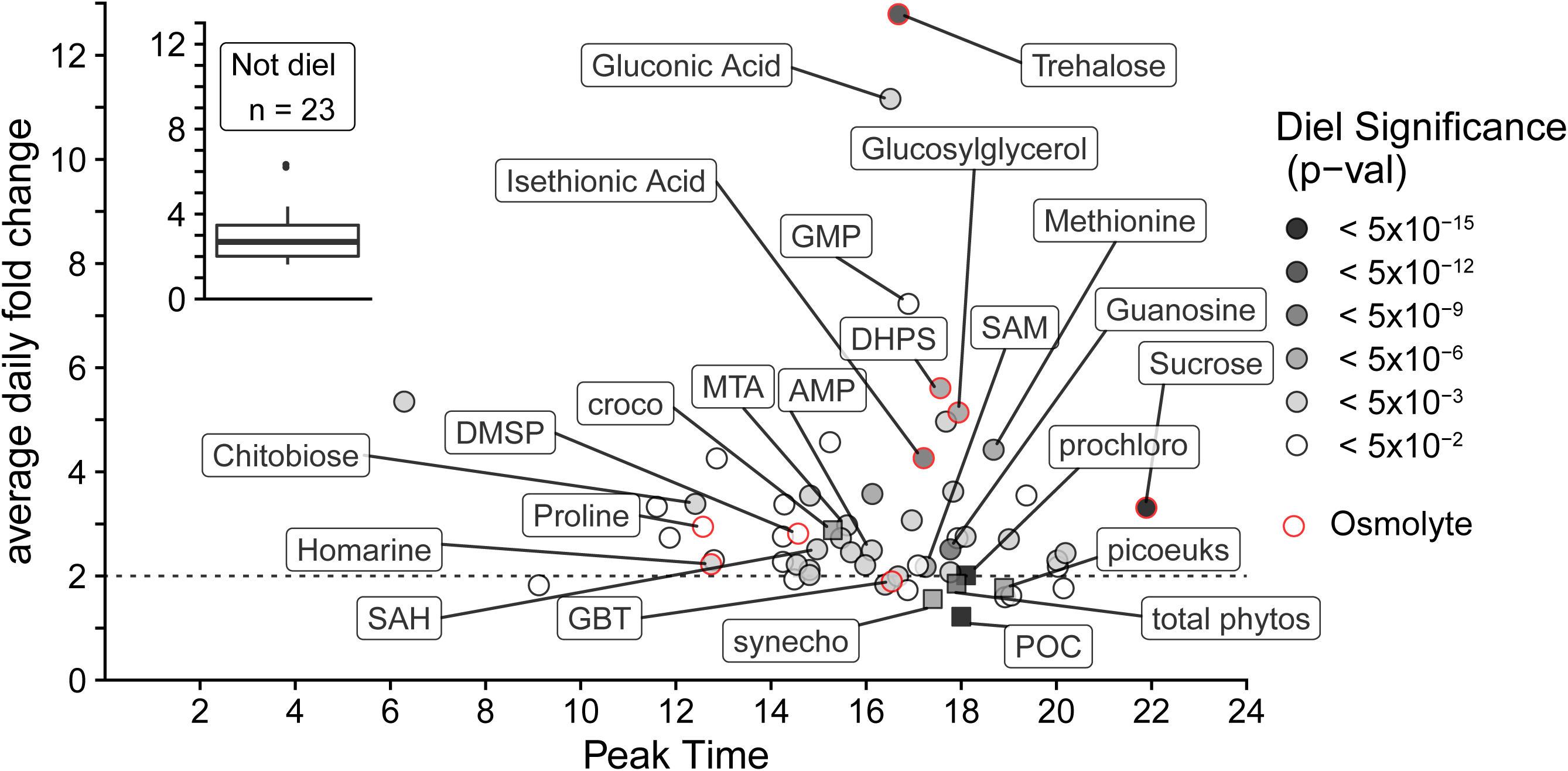
Peak time vs average daily fold change for each metabolite (circles, nmol L-1), POC from beam attenuation and phytoplankton biomass from flow cytometry (squares, µg C L-1). Grey color indicates the level of significance (fdr corrected p-value) of the 24-hour oscillation. The inset shows the distribution of fold-change in non-significant compounds. These compounds have variability even though they do not have 24-hour periodicity. Red outlines indicate that the compound is an osmolyte. Select compounds and all biomass estimates are labeled (croco = Crocosphaera, synecho = Synechococcus, prochloro = Prochlorococcus, picoeuks = picoeu-karyotes, total phytos = total phytoplankton biomass from underway flow cytometry).

**Figure 5.**
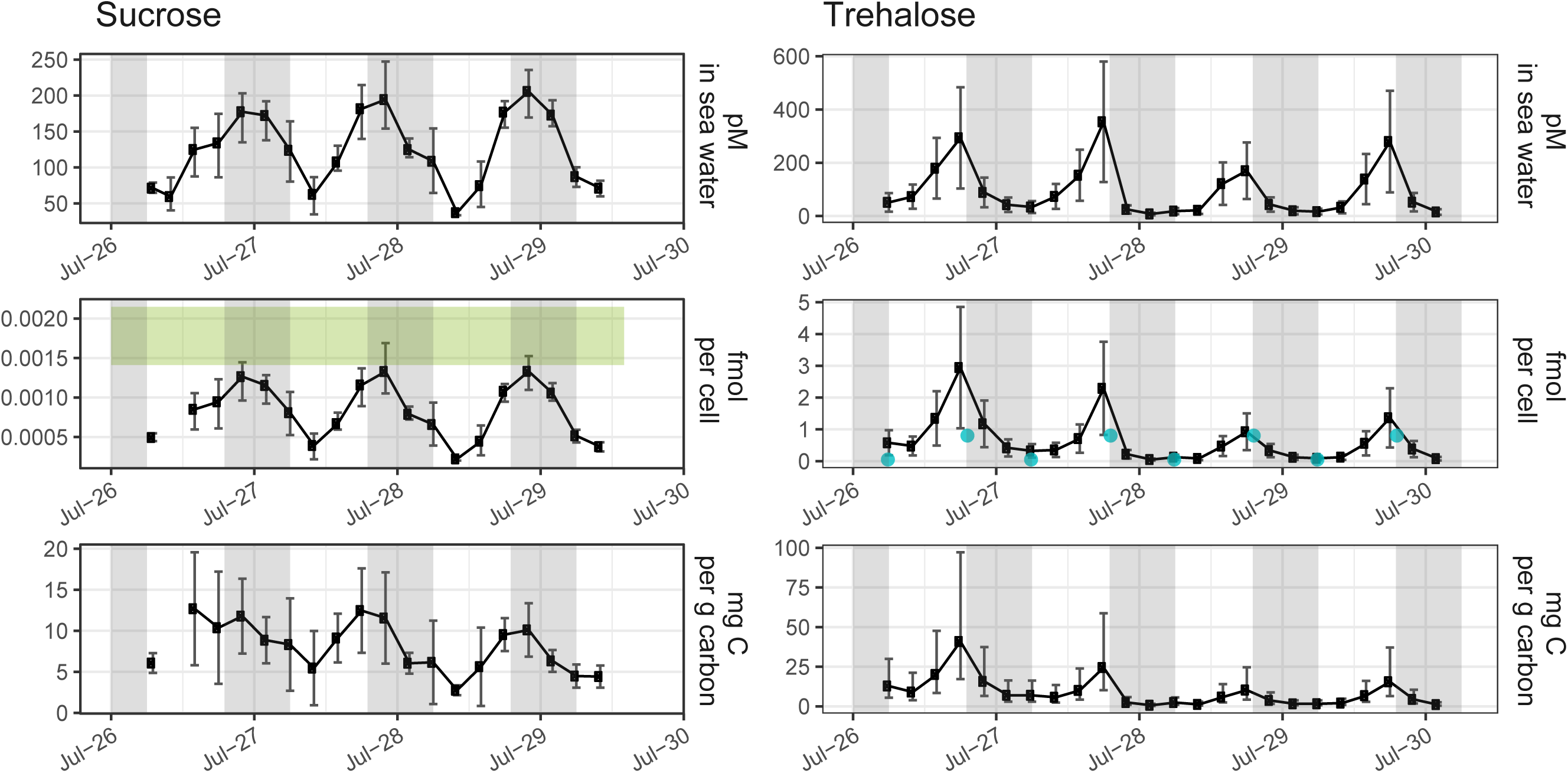
Particulate sucrose (left) and trehalose (right) measured as pmol L-1 in seawater (top), fmol cell-1 (middle) of Crocosphaera and Procholorococcus for trehalose and sucrose, respectively, and mg g-1 cell carbon (bottom) of Crocosphaera and Procholorococcus for trehalose and sucrose, respectively. The light grey vertical shading represents nighttime. The green box in the middle-left panel indicates the range of cellular sucrose quotas measured in Procholorococcus MIT1314 harvested mid-day in exponential growth. The blue points in the middle-right panel indicate the dawn and dusk values measured for trehalose quotas in Crocosphaera watsonii strain WH8501. In the top panels, the error bars represent one standard deviation around the mean value, including uncertainty from the quantification regression. The error bars in the middle panels represent one standard deviation around the mean. The error bars in the bottom panels represent the 95% confidence interval given the confidence in the biomass quantification from underway flow cytometry.

**Figure 6:**
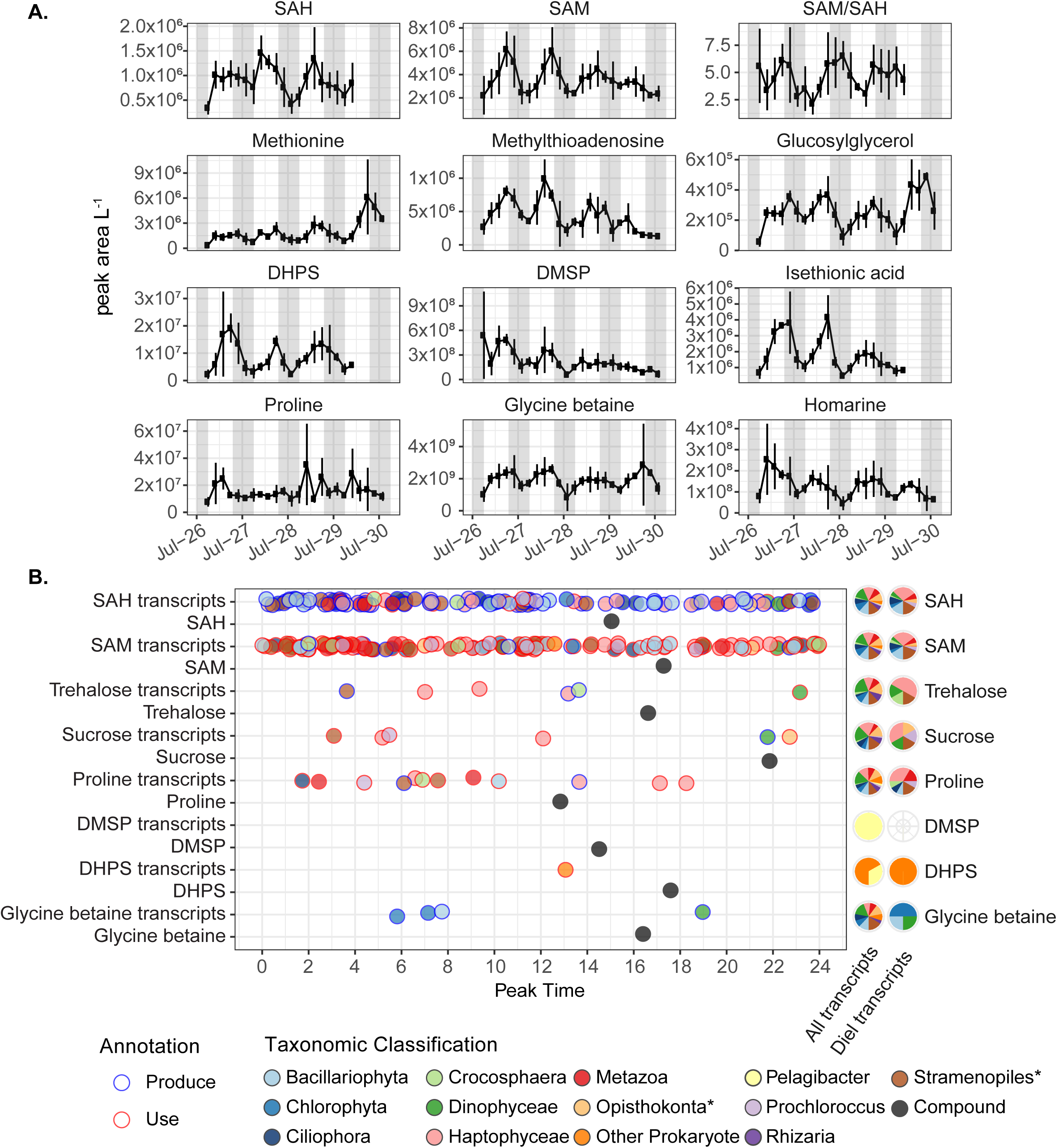
A) Diel metabolite concentrations (peak area L-1, proportional to nmol L-1) of methionine cycle compounds, methylth-ioadenosine, and osmolytes. Error bars are the standard deviation of biological triplicates. The light grey vertical shading represents nighttime. B) Left: Time of peak abundance of diel transcripts related to the production or use of select diel osmolytes and primary metabolites. Fill color indicates the phylogenetic lineage of the transcript; outline color indicates whether the transcript is associated with production or consumption of the metabolite. Time of metabolite peak concentration (nmol L-1) is in black. Right: Proportion of all transcripts and diel transcripts belonging to each taxon. * = does not include select subgroups shown otherwise.

Primary metabolites involved in anabolism and redox balance showed diel oscillations. The three methionine-cycle compounds detected, S-adenosyl methionine (SAM), S-adenosyl homocysteine (SAH), and methionine, showed oscillations. 5’-Methylthioadenosine (MTA) is produced from SAM during polyamine synthesis and had a temporal pattern that closely matched SAM (Figures 3, 6), such that SAM/MTA remained relatively constant. Pantothenate (Vitamin B_5_) was one of the few compounds that peaked in the morning (Figure 3). Vitamins involved in redox balance, riboflavin and niacin (Vitamins B_2_ and B_3_) oscillated with maxima near dusk. Reduced glutathione oscillated with an afternoon peak (Figure 3).

### Connections between metabolites and transcripts

To investigate the relationships between gene expression and metabolite concentration we used the KEGG database to connect metabolites with transcripts annotated as encoding proteins that directly produce or degrade each metabolite. All but four of our diel metabolites related to at least one annotated prokaryotic or eukaryotic transcript (Figure S2). Glucosylglycerol, ergosterol, and isethionic acid are in the KEGG database but no transcripts were annotated in our dataset as directly producing or degrading them, while homarine is not included in the KEGG database.

Although the number of transcripts associated with each metabolite is inherently biased by the databases used and the depth of sequencing, transcripts provide insight into the number and identity of organisms and pathways that may be responsible for the metabolite’s synthesis and degradation. The orders containing *Crocosphaera, Prochlorococcus, Pelagibacter ubique*, and other unclassified alphaproteobacteria comprised ∼50% of all prokaryotic transcripts that could be linked to metabolites (Table S6).. Dinoflagellates (Dinophyceae), non-diatom stramenopiles (Stramenopiles), haptophytes (Haptophyceae), non-metazoa opistokonts (Opisthokonta), and diatoms (Bacillariophyta) comprised ∼70% of eukaryotic transcripts linked with metabolites (Table S7). Adenosine monophosphate (AMP), SAM, and SAH stand out as the diel metabolites with the largest number of associated diel transcripts, with 181, 124, and 113 transcripts respectively (Figure 6, Figure S2). Most diel SAM and SAH transcripts were methyltransferases that convert SAM into SAH (Tables S6, S7). In most other cases, there were few diel transcripts associated with a metabolite (e.g. only 6 diel genes were associated with trehalose, Figure 6, Figure S2).

To investigate the temporal relationship between gene expression and metabolite concentration, we estimated the lag-time between metabolites and transcripts that exhibited significant diel periodicity. This analysis showed a broad distribution in the lag-times between metabolites and transcripts, with no predictable lag for prokaryotic or eukaryotic transcripts and their associated metabolites. (Figure S3).

### Disaccharide osmolytes can be attributed to cyanobacteria

We observed trehalose-related transcripts from eukaryotic phytoplankton and *Crocosphaera* (Figure 6). Using published *Ostreococcus* cellular trehalose concentrations(20) and picoeukaryote cell counts, we estimated that picoeukaryote contribution to trehalose was 0.2–3.0 pmol L^-1^, a small fraction of environmental trehalose (up to 627 pmol L^-1^). The abundance of *Crocosphaera* (Table 1, Figure 1) and diel oscillations in the *Crocosphaera* transcript for trehalose 6-phosphate synthase/phosphatase (Figure 6, Table S6) suggest *Crocosphaera* as the main contributor of trehalose during this field study. To test this hypothesis, we grew *Crocosphaera watsonii* WH8501 under a 12:12 light:dark cycle and measured 0.8 and 0.07 fmol trehalose cell^-1^ at the end of the light and dark periods, respectively (Figure 5, Figure S4). Given the *Crocosphaera* abundance during our sampling and assuming similar intracellular concentration, this accounts for 1.8–670 pM particulate trehalose, comparable to total particulate trehalose during our sampling (2.8–627 pmol L^-1^ across both sampling periods, Figure 5).

Multiple taxa expressed transcripts related to production and degradation of sucrose, including *Prochlorococcus* (Figure 6). To assess the potential contribution of *Prochlorococcus* to environmental sucrose concentrations, we measured the cellular sucrose quota in a culture of *Prochlorococcus* MIT1314 harvested midday during exponential growth. Using the cellular quota of sucrose in these cultures (range in biological triplicates: 1.4–2.1 amol cell^-1^) and the abundance of *Prochlorococcus* at the time of sampling, it is possible that all the observed sucrose could have been in *Prochlorococcus* during this study (Figure 5).

## Discussion

As a whole, the metabolites we measured comprise up to 2% of POC and 3% of PN in our samples (Figure 2B,C). This is a reasonable value given ∼80% of surface POC is comprised of lipid, carbohydrate, and protein macromolecules(16,17), and DNA, RNA, and pigments contribute several percent of the dry weight of actively growing microalgae(55). Metabolite pools are dynamic, and an increase in the concentration of a given metabolite suggests that sources of that compound (biosynthesis, uptake from dissolved pools, or polymer disassembly) are greater than sinks (exudation, loss due to cell death, intracellular degradation, or polymer assembly). The prevalence and amplitude of diel oscillations in metabolite concentrations reflect that many members of the surface microbial community near Station ALOHA were synchronized to diel light periodicity.

### Community synchrony is driven by diel partitioning of anabolism, catabolism, and redox maintenance

The diel oscillations in POC and FCM-resolvable phytoplankton biomass reflect the alternation of carbon fixation, anabolism, and growth during daylight hours and respiration, catabolism, and mortality during the night (Figure 1)(2,3,5). The community metabolome reflects these patterns with an overall increase in concentration throughout the day and a consistent morning phenotype (Figure 2, Figure S1A,B), reflecting nighttime use of energy stores and recovery from daytime oxidative stress(56). Nearly half of the diel metabolites (26/55) had peak molar concentrations near dusk (Figure 3), corresponding with a peak in carbon biomass. However, for most (46/55) diel metabolites, the daily enrichment of a metabolite exceeded that of POC or total FCM-resolvable phytoplankton biomass, which had daily fold changes of 1.2 and 1.8, respectively (Figure 4). This suggests these metabolites likely had oscillations in intracellular concentration, as previously observed for many primary metabolites in non-marine cyanobacteria(57).

Primary metabolites are particularly powerful indicators of biochemical activity on the community scale. SAM, SAH, and AMP are compounds involved in biosynthesis that had diel oscillations with daytime increases (Figures 3, 6). Individual transcripts associated with these molecules had diel patterns that peaked at all times of day, across a myriad of pathways and microbial taxa (Figure 6, Figure S2). Despite this diversity in use, the sum of community activity was reflected in diel oscillations of metabolite concentrations, which were synchronized with daytime biomass accumulation. Further evidence of this daytime community-scale anabolism is the diel oscillation of pantothenate (Vitamin B_5_), a component of Coenzyme A as well as Acyl Carrier Protein. Pantothenate peaked in the morning (Figure 3), suggesting that the community was poised to assemble these cofactors for daytime biosynthesis.

SAM is a ubiquitous methyl donor used by all living cells. During methylation, SAM is converted to SAH, which is then regenerated back to SAM via methionine. In addition to its role in methylation, SAM is essential for polyamine synthesis and is the most common riboswitch effector in prokaryotes(58). SAM riboswitches have been observed in native Station ALOHA bacterioplankton populations(59). SAH had an afternoon peak time, such that the SAM/SAH ratio was at a minimum during the day (Figure 6). This ratio reflects methylation potential, suggesting that the demand for methylation outstripped the supply of SAM in the light. Over the dark period, SAM/SAH ratios recovered, suggesting that catabolic processes dominated and the need for SAM was diminished. Many cells require cobalamin to catalyze the reactions that regenerate methionine, and SAH is elevated relative to SAM during cobalamin stress as cells struggle to complete the cycle(25). Thus, it is possible that the lower SAM/SAH ratio additionally reflects a daytime increase in cobalamin usage.

Managing oxidative stress is a critical part of cellular activity. Reactive oxygen species produced during photosynthesis accumulate over the day and present a continuing challenge for cells at night(60). Riboflavin and niacin (vitamins B_2_ and B_3_) are involved in redox balance and show similar daytime accumulations (Figure 3, Table S3). These are precursors to FMN/FAD and NAD/NADP, respectively, and reflect community-wide diel patterns of redox processes. Cyanobacteria manage excess energy during the day by storing glycogen and producing small molecules that can either be stored or excreted(56,60–63). At night, glycogen is catabolized via hydrolysis followed by glycolysis or the oxidative pentose phosphate pathway (OPPP), producing the reductant sources NADH and NADPH. Gluconic acid accumulation during the day (Figures 3, 4) may reflect less flux through OPPP during the day, while photosynthesis produces NADPH, followed by a switch towards OPPP at night(56). Reduced glutathione also showed a daytime peak (Figure 3), as has been observed in cultures and field studies(64), possibly reflecting production to compensate for increased oxidative stress in the day, and a subsequent decrease in production and oxidation of the residual pool overnight.

### Diel oscillations in osmolyte concentrations reveal complexity in their function

Metabolites with osmolyte properties are among the most abundant compounds within marine microbial cells(27,65–69) and exhibited diel oscillations (Figures 5, 6). One exception to this observation was glutamic acid, which plays a critical role in regulating nitrogen assimilation in addition to its osmotic properties(65). In the absence of fluctuations in salinity or temperature, oscillations in osmolyte concentrations occurred in excess of or out of sync with biomass oscillations and point to alternative roles for this compound group(65) (Figure 4, Table S3). Intracellular accumulation of osmolytes occurred predominantly during the day when electron flow through the photosystems and the Calvin Cycle exceeds that required to maintain maximum division rates. The resulting need to dissipate reductant is typically channeled into the production of carbohydrates like glycogen(8,56,63), exopolymeric substances(70,71), or into storage lipids(12,72). These energy stores are used to fuel cellular respiration and other activities at night, such as protein synthesis and preparing cells for photosynthesis(12,56,63,72). Unlike starch and storage lipids, osmolytes do not necessarily need to go through hydrolysis, β-oxidation, or glycolysis prior to entering the TCA cycle, and could be used as readily available substrates for energy production and as biosynthetic intermediates while macromolecular pools are being mobilized by the cell(61).

Trehalose was the most prominent diurnally oscillating compound (Figures 4, 5). Trehalose is an osmolyte produced by the unicellular diazotroph *Crocosphaera*(68,73), some heterotrophic bacteria, and some phytoplanktonic picoeukaryotes, including *Ostreococcus*(20). Transcriptomic evidence motivated us to measure trehalose in cultures of *Crocosphaera*, which revealed differences in intracellular trehalose at the beginning and end of the day. Assuming trehalose in the environment is produced primarily by *Crocosphaera*, our results strongly suggest that intracellular trehalose concentrations have diel oscillations in the field (Figure 5).

*Crocosphaera* temporally separate photosynthesis and nitrogen fixation to protect nitrogenase from oxygen(74–76), they therefore need energy at night to draw down cellular oxygen and fuel nitrogen fixation(77,78). *Crocosphaera* has at least one gene encoding a protein homologous to glycoside hydrolases, family 15(79), which contains enzymes that hydrolyze a variety of glycosidic bonds, including trehalose. Thus, it is possible that *Crocosphaera* use trehalose as a fuel for generating the electrons and ATP required for nitrogen fixation. Using the stoichiometry of these reactions(77,80), we estimated that trehalose catabolism could have fueled 9–28% of the nighttime nitrogen fixation during this expedition(7) (calculation in supplemental material). As much as 60% of total dark respiration by *Crocosphaera* is used to draw down cellular oxygen rather than to directly fuel nitrogen fixation(77), and, if we adjust our calculation accordingly, trehalose can produce 3.6–11% of the required respiratory substrates needed for *Crocosphaera* to effectively fix nitrogen at the rates measured(7).

The flux of carbon through trehalose may be an indicator of the accumulation and degradation of a larger glycogen pool that accumulates during the day and is used at night(81). Shi et al. (2010) suggest that *Crocosphaera* cells are depleted of storage compounds at night, since prolonged dark does not result in increased nitrogen fixation(82). If this hypothesis is correct, the total amount of nitrogen fixation possible is limited by the amount of energy stored in substrates such as trehalose and glycogen during daytime, and the ability to accumulate and use these compounds could have impacts on the nitrogen budget of the microbial community.

Another disaccharide osmolyte, sucrose, displayed an oscillation with a maximum daily concentration at 22:00. Sucrose is the major compatible solute in high-light *Prochlorococcus* (67), and the observed environmental variation may reflect the *in situ* accumulation and use of glycogen by *Prochlorococcus*. Though other organisms also expressed sucrose related genes (Figure 6), *Prochlorococcus* was the numerically dominant sucrose-producing organism detected in these populations (Table 1) and is known to accumulate polysaccharides during the day, particularly under nitrogen limitation(83). If we assume that cellular quotas of sucrose in *Prochlorococcus* grown in culture are like those in the environment, *Prochlorococcus* alone could explain the sucrose concentrations seen in the environment (Figure 5). Sucrose had a diel oscillation when normalized to *Prochlorococcus* cell counts and biomass (Figure 5, Figure S5). These potential intracellular oscillations lead us to hypothesize that *Prochlorococcus* uses sucrose for energy storage and not only as a compatible solute, as has been observed in non-marine cyanobacteria(57,61).

Homarine and DMSP are known eukaryotic osmolytes(65,66,69,84). Here the amplitude and timing of the diel oscillations in these two compounds differ from those observed in phytoplankton picoeukaryote biomass (Figure 4), suggesting that these compatible solutes play additional roles within the microbial community. This diversity of functions is well established for DMSP, which influences grazing behaviors and can function as an antioxidant(69,85). DMSP is also a carbon and reduced sulfur source in the microbial community, with uptake and assimilation both tied to light availability(86,87). In our analysis, the only transcript related to DMSP encodes a SAR11 DMSP demethylase required for DMSP degradation (Figure 6). A dearth of data on the roles of homarine in marine microbes and a lack of genetic information about homarine synthesis and degradation limit our ability to infer the sources and sinks for this abundant compound. The high concentration and diel dynamics of homarine calls for further investigation.

Both isethionic acid and DHPS are associated with fast growing eukaryotes that need to mobilize cellular machinery to transport materials into the mitochondria for respiration(27,88), and recent work has suggested that DHPS has potential osmotic capabilities(27). These two metabolites had large diel oscillations implicating them as temporary stores of energy or intermediates that can be mobilized quickly. Our data implicates SAR11 and Rhodobacteraceae as likely DHPS degraders at Station ALOHA (Figure 6), although genes for the production of DHPS are not in the KEGG database and thus were not identified by our analyses. If production and degradation of these compounds are separated along phylogenic lines(31) then these compounds are likely excreted into the dissolved phase by eukaryotes and subsequently available for use by bacteria, as suggested in Durham et al. (2019). This may explain the midday maximal expression of a *hpsN*-like Rhodobacteraceae DHPS degradation gene (Figure 6).

### Metabolites as fuel for the microbial loop

Several of the metabolites investigated here are known to fuel heterotrophic bacterial growth in marine ecosystems(87,89–92). DMSP, for example, can support up to 9.5% of the bacterial carbon demand at Station ALOHA(86). Particulate metabolite concentrations and their oscillations observed in this study call for further investigation into the hypothesis that these compounds are important substrates for community interactions and resources for the microbial loop. For compounds that exhibited diel oscillations, the difference between the daily maximum and minimum values provides a daily net production and degradation rate. We estimated a total net turnover rate of over 27 nmol C L^-1^ d^-1^ from our targeted metabolites, with several metabolites exhibiting individual turnover rates of over 1 nmol C L^-1^ d^-1^, including arachidonic acid, trehalose, homarine, sucrose, GBT, glucosylglycerol, and DHPS (Table S3). These are conservative estimates since the instantaneous flux may be much higher than the daily net change and we did not measure excretion of metabolites into the dissolved pool. For example, DMSP has a turnover time of 4.5 hours at Station ALOHA(86) and has been shown to be produced at night and during the day(93), both observations would substantially increase the baseline estimate of DMSP production which does not account for rapid turnover and only includes a daytime increase in concentration. While the fate of the metabolites measured here remain unclear, conservative estimates of carbon and nitrogen flux through these small pools was large, comprising around 2% of the ^14^C based estimates of primary productivity during this study(12). These compounds are potentially used for cellular requirements by the organisms synthesizing them, as discussed above, or released into the labile dissolved pool. When they enter the dissolved pool through excretion or cell lysis, these compounds are important components of the labile dissolved organic matter pool(91) and play a role in organism interactions(94,95).

## Conclusions

The light-dark cycle plays a dominant role in structuring marine microbial activity. Previous work has shown diel oscillations of community processes, such as daily accumulation and depletion of POC(2), and diel oscillations of transcriptional activity, which have provided new information on temporal dynamics and raise hypotheses about the activity of individual taxa(9,10). Measurements of *in situ* metabolites in native planktonic microbial populations reported here support the hypotheses that diverse microbial taxa in the NPSG are synchronized to daily oscillations of light energy and photosynthesis, with metabolites accumulated during the day and depleted at night. The diel synchrony of ubiquitously used primary metabolites shows the extent to which photoautotrophic organisms dominate the community and drive anabolic processes during the day and catabolic processes at night. The combination of transcript abundances, metabolite concentrations, and taxa-specific biomass in the field and in culture allows us to postulate that *Crocosphaera* uses trehalose as a short-term energy source to drive nighttime nitrogen fixation. Trehalose and the other osmolytes we measured are highly abundant in cells and, in addition to playing multiple roles in their producers, likely fuel the metabolism of heterotrophic bacteria. Metabolite concentrations cannot be predicted from transcripts in a single organism in pure culture, let alone in a complex natural community. Pairing quantitative measurements of particulate metabolites with transcriptomes is a key step toward understanding how regularly oscillating gene expression in microbial communities is reflected in the net community processes we observe and further elucidates the currencies of the microbial community.

## Acknowledgements

The authors acknowledge A. Hynes, N. Kellogg, R. Lionheart, M. Motukuri, and A. Wied for assistance with lab and data analysis; J.S. Weitz and D. Muratore for productive discussions and feedback; A. E. White for the POC and PN data; J.P. Zehr and M. Hogan for providing *Crocosphaera WH8501*; A. Coe and S.W. Chisholm for providing *Prochlorococcus* MIT1314; the crew and scientific party of the R/V *Kilo Moana* during HOE-Legacy 2A. This work was supported by grants from the Simons Foundation (LS Award ID: 385428, A.E.I.; SCOPE Award ID 329108, A.E.I., E.F.D., E.V.A.; SCOPE Award ID 426570, E.V.A.; Award ID 598819, K.R.H.), the National Science Foundation (NSF OCE-1228770 and OCE-1205232 to A.E.I., NSF OCE-160019 to R.D.G, NSF GRFP to A.K.B. and K.R.H., NSF IGERT Program on Ocean Change to A.K.B), and the Gordon and Betty Moore Foundation (Grant #3777 to E.F.D.)

## Competing interests

The authors declare no conflict of interest.

## Data availability

Information for the KM1513/HOE Legacy II cruise can be found online at http://hahana.soest.hawaii.edu/hoelegacy/hoelegacy.html. Raw sequence data for the diel eukaryotic metatranscriptomes are available in the NCBI Sequence Read Archive under BioProject ID PRJNA492142. Raw sequence data for the prokaryotic metatranscriptomes are available in the NCBI Sequence Read Archive under BioProject ID PRJNA358725. Raw and processed metabolomics data are available in Metabolomics Workbench under Project ID PR000926, currently embargoed until July 1, 2020 but available upon publication.

## Author contributions

AEI, EVA, and EFD designed the study. AKB, LTC, FOA, BPD, and FR collected the samples. AKB, LTC, and KRH performed the metabolite sample processing and analysis. FOA, RDG, and BPD performed the metatranscriptomic sample processing and analysis. FR performed the flow cytometry sample processing and analyses. AKB, KRH, and AEI contributed to data interpretation and visualization. AKB and AEI drafted the paper and incorporated revisions from all authors.

---

Supplemental Figure 1

Multivariate analyses based on *z-*scored particulate metabolite concentration (proportional to nmol L^-1^). A) NMDS of the first sampling period alone: Jul-26^th^ - Jul 30th. The NMDS analysis results were significant (Monte Carlo randomization *p* < 0.01) with a stress value of 0.18. B) Within and between group variability from ANOSIM analysis using *z*-score standardized particulate concentrations of all metabolites (nmol L^-1^) from the first sampling period (*R* = 0.194, *p* < 0.001). C) NMDS of the second sampling period alone: Jul 31st – Aug 3rd. The NMDS analysis results were significant (Monte Carlo randomization *p* < 0.01) with a stress value of 0.17. D) NMDS of full dataset: Jul-26th – Aug-3rd. Colors indicate time of day that the samples were collected. The NMDS analysis results were significant (Monte Carlo randomization *p* < 0.01) with a stress value of 0.18.

Supplemental Figure 2

Diel transcript peak abundance related to the production or degradation of diel metabolites. Color indicates the phylogenetic lineage of the transcript. Left: Peak time of transcript abundance or particulate metabolite concentration (nmol L^-1^). Right: Proportion of diel transcripts belonging to each taxa and proportion of all transcripts, regardless of diel oscillation, related to each metabolite belonging to each taxa.

Supplemental Figure 3

Offset time (in hours) between the diel compounds and diel eukaryotic transcripts (top) or diel prokaryotic transcripts (bottom) that use or produce them. Diel significance of compounds was based on the first sampling period, diel significance of eukaryotic transcripts was based on the first sampling period, diel significance of the prokaryotic transcripts was based on both sampling periods (RAIN fdr-corrected *p* < 0.05).

Supplemental Figure 4

Field and culture particulate trehalose concentrations normalized to *Crocosphaera* cell count. Field data (black points) show the average and standard deviation at each time point over the full sampling period. Lab cultures (green circles) represent the values for the cultures harvested at dawn and dusk. Variability in technical replicates (for dusk) and biological duplicates (for dawn) are smaller than the points.

Supplemental Figure 5

Field and culture sucrose per cell *Prochlorococcus*. Field data (black points) show the median and range at each time point. The green box shows the maximum and minimum values of sucrose in triplicate axenic cultures of *Prochlorococcus* MIT1314 harvested at mid-day in exponential growth.

Supplemental Figure 6

Time of day that compounds peak in the second sampling period.

Supplemental Table S1

Internal Standards added before extraction (Exr Standard) or before injection (Inj Standard)

Supplemental Table S2

Transcriptomes used to supplement the MarineRefII reference database (http://roseobase.org/data/).

Supplemental Table S3

Metabolites measured in this analysis. The average fold change from peak to trough, the maximum and minimum estimated or absolutely quantified values (pmol L-1), and whether the compound oscillates with 24-hour periodicity when normalized to volume of seawater filtered (water), when normalized to POC (POC), both (Both), or neither (None) for the first sampling period analyzed independently, second sampling period analyzed independently, and full dataset. The time of peak concentration for these various normalizations and time periods are provided in the final columns, rounded to the nearest hour. The net flux through the particulate pool calculated by the mean daily swing from max to minimum. * indicates metabolites for which samples 21-24 are removed and for which 6 samples in the second diel sampling period maybe affected by internal standard adjustments. + indicates metabolites for which 4 samples in the second sampling period might affected by IS adjustments. † notes that concentrations for DMSP are likely underestimates, as described in the methods.

Supplemental Table S4

Average and standard deviation of targeted metabolite composition at dawn (06:00) and dusk (18:00) from July 26^th^ – July 28^th^ (*n* = 9 for each time point), as the estimated particulate metabolite concentration, the percent of particulate organic carbon, and the percent of the particulate nitrogen.

Supplemental Table S5

Pairwise comparisons of samples collected at different time points from the multivariate analyses of particulate metabolite concentration during the first sampling period.

Supplemental Table S6

Prokaryotic transcripts that matched metabolites identified by organism taxa and KEGG ortholog. If the transcript is significantly diel (RAIN fdr-corrected *p*-value < 0.05) the time of peak transcript abundance is provided (0/24 is midnight, 12 is noon).

Supplemental Table S7

Eukaryotic transcripts that matched metabolites identified by organism taxa and KEGG ortholog. If the transcript is significantly diel (RAIN fdr-corrected *p*-value < 0.05) the time of peak transcript abundance is provided (0/24 is midnight, 12 is noon).

Supplemental Table S8

Particulate metabolite concentrations in normalized peak area per L of seawater filtered. Across a single metabolite these values are proportional to molar concentration. Values should not be quantitatively compared between two metabolites, since the ionization efficiency and matrix effects influence different metabolites differently such that the same concentration can result in difference in peak area.

## Notes

### Competing Interest Statement

The authors have declared no competing interest.

