## Supplemental Figures for "Diel Oscillations of Particulate Metabolites Reflect Synchronized Microbial Activity in the North Pacific Subtropical Gyre"

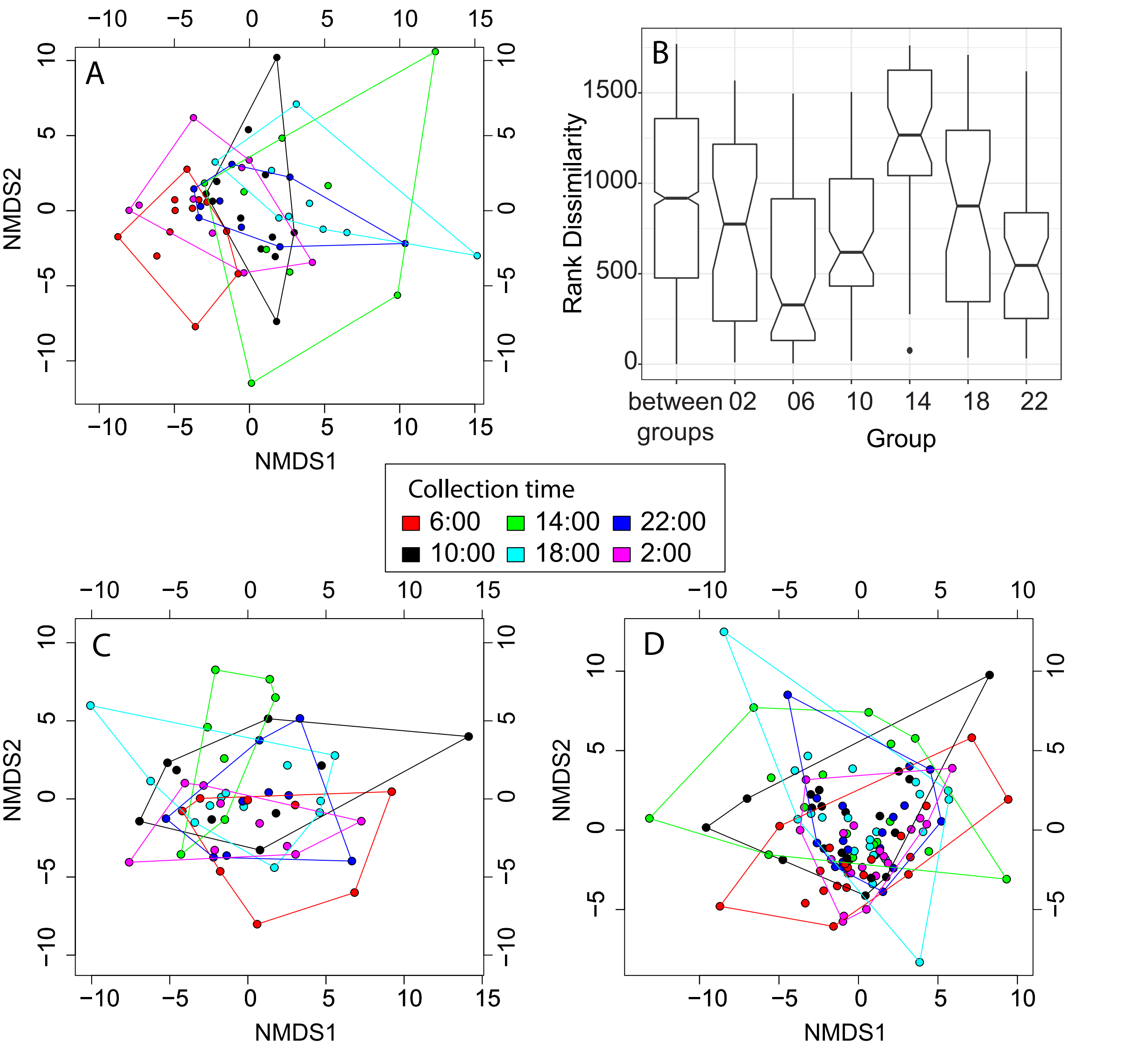


Supplemental Figure 1. Multivariate analyses based on z-scored particulate metabolite concentration (proportional to nmol L-1). A) NMDS of the first sampling period alone: Jul-26th - Jul 30th. The NMDS analysis results were significant (monte carlo randomization $p < 0.01$) with a stress value of 0.18. B) Within and between group variability from ANOSIM analysis using z-score standardized particulate concentrations of all metabolites (nmol L-1) from the first sampling period ($R = 0.194$, $p < 0.001$). C) NMDS of the second sampling period alone: Jul 31st – Aug 3rd. The NMDS analysis results were significant (monte carlo randomization $p < 0.01$) with a stress value of 0.17. D) NMDS of full dataset: Jul-26th – Aug-3rd. Colors indicate time of day that the samples were collected. The NMDS analysis results were significant (monte carlo randomization $p < 0.01$) with a stress value of 0.18.

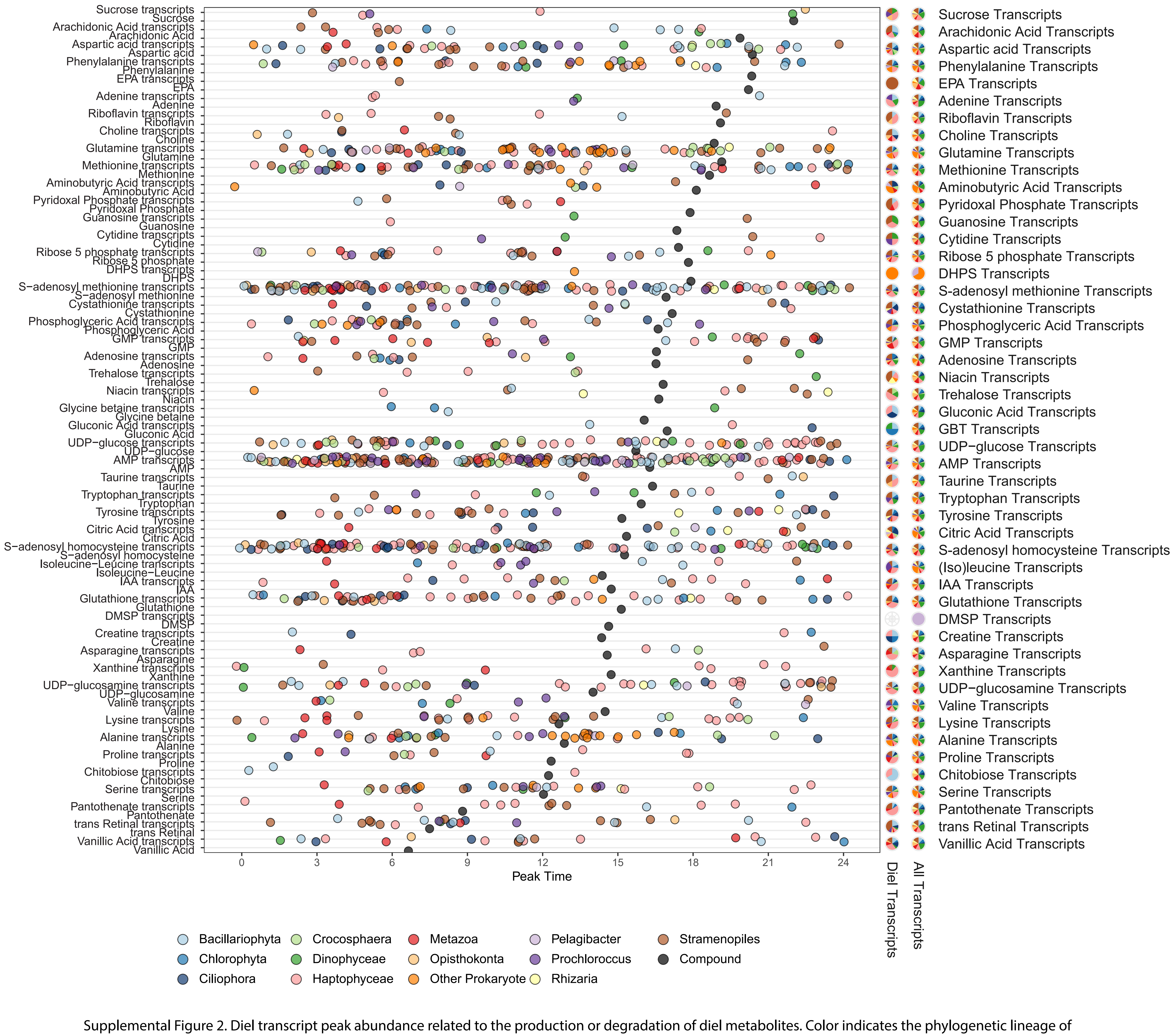


Supplemental Figure 2. Diel transcript peak abundance related to the production or degradation of diel metabolites. Color indicates the phylogenetic lineage of the transcript. Left: Peak time of transcript abundance or particulate metabolite concentration (nmol L-1). Right: Proportion of diel transcripts belonging to each taxa and proportion of all transcripts, regardless of diel oscillation, related to each metabolite belonging to each taxa.

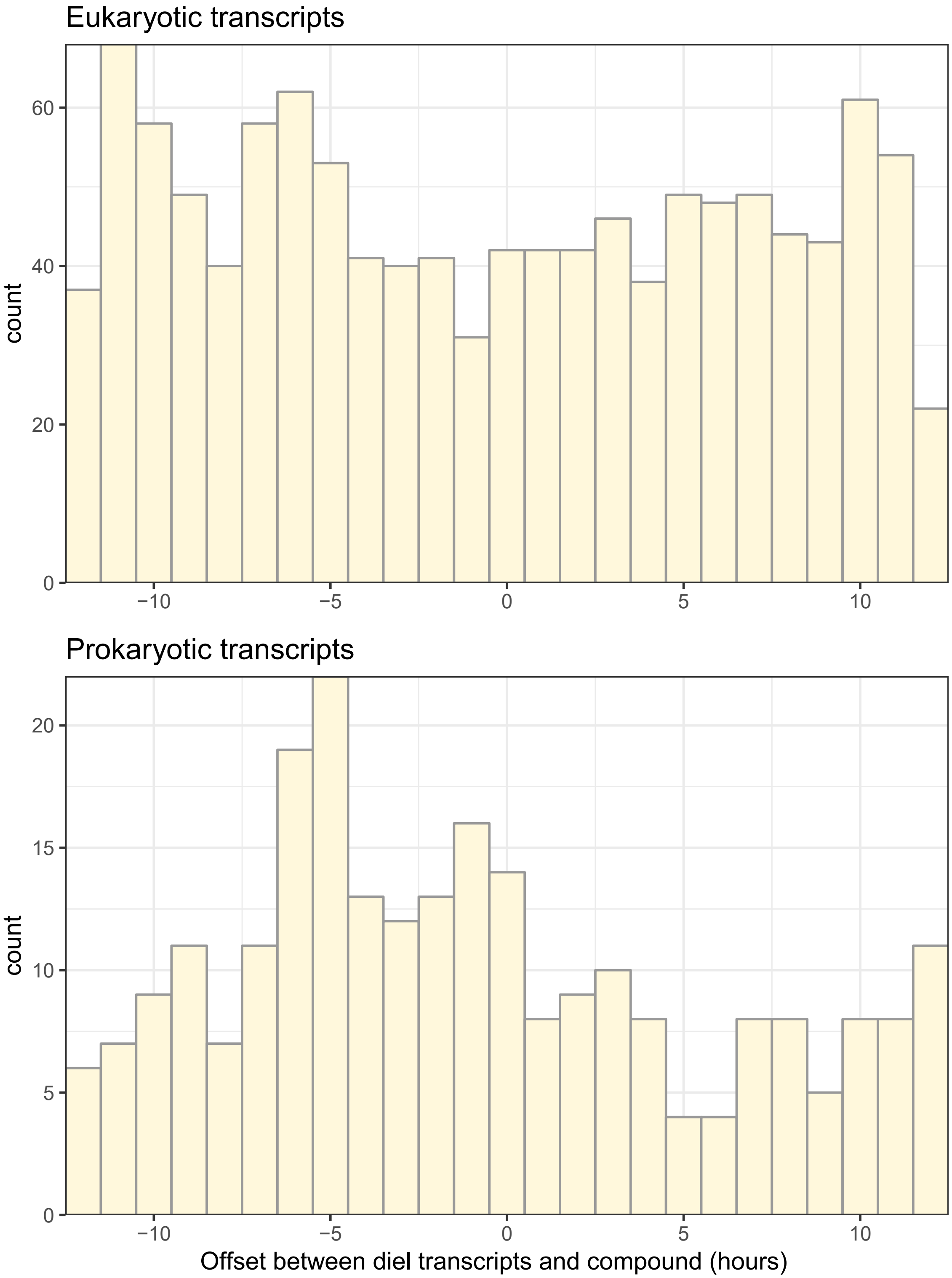


Supplemental Figure 3. Offset time (in hours) between the diel compounds and diel eukaryotic transcripts (top) or diel prokaryotic transcripts (bottom) that use or produce them. Diel significance of compounds was based on the first sampling period, diel significance of eukaryotic transcripts was based on the first sampling period, diel significance of the prokaryotic transcripts was based on both sampling periods (RAIN fdr-corrected $p < 0.05$).

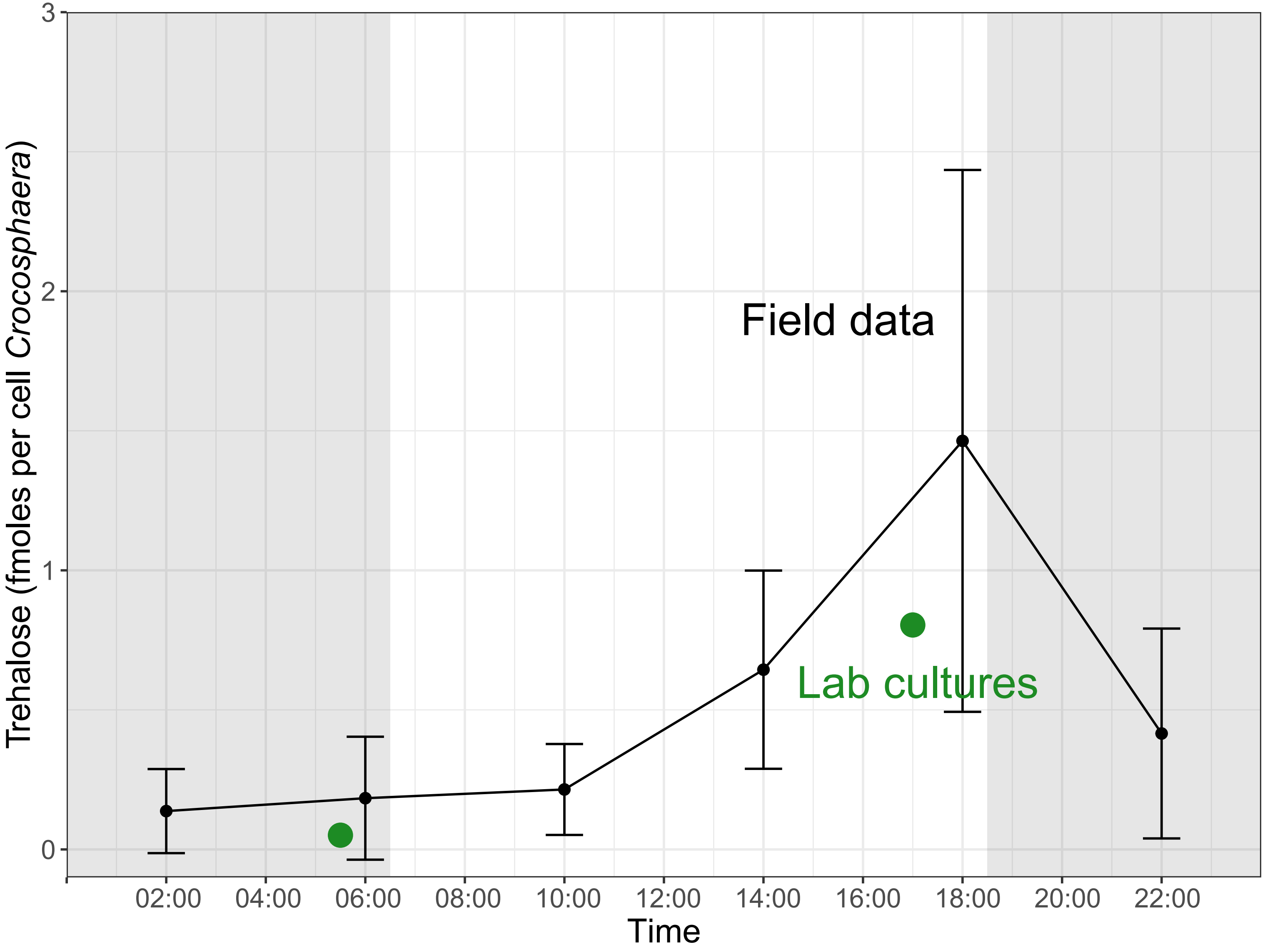


Supplemental Figure 4. Field and culture particulate trehalose concentrations normalized to Crocosphaera cell count. Field data (black points) show the average and standard deviation at each time point over the full sampling period. Lab cultures (green circles) represent the values for the cultures harvested at dawn and dusk. Variability in technical replicates (for dusk) and biological duplicates (for dawn) are smaller than the points.

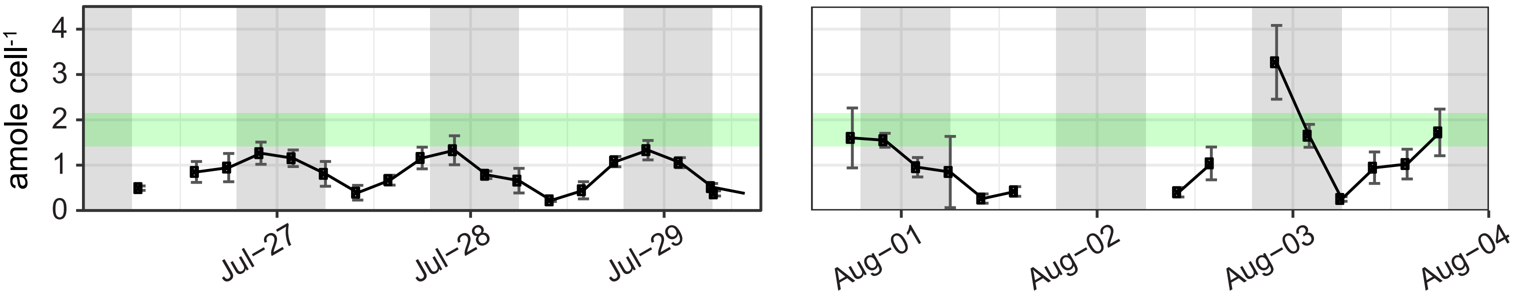


Figure S5. Field and culture sucrose per cell Prochlorococcus. Field data (black points) show the median and range at each time point. The green box shows the maximum and minimum values of sucrose in triplicate axenic cultures of Prochlorococcus MIT1314 harvested at mid-day in exponential growth.

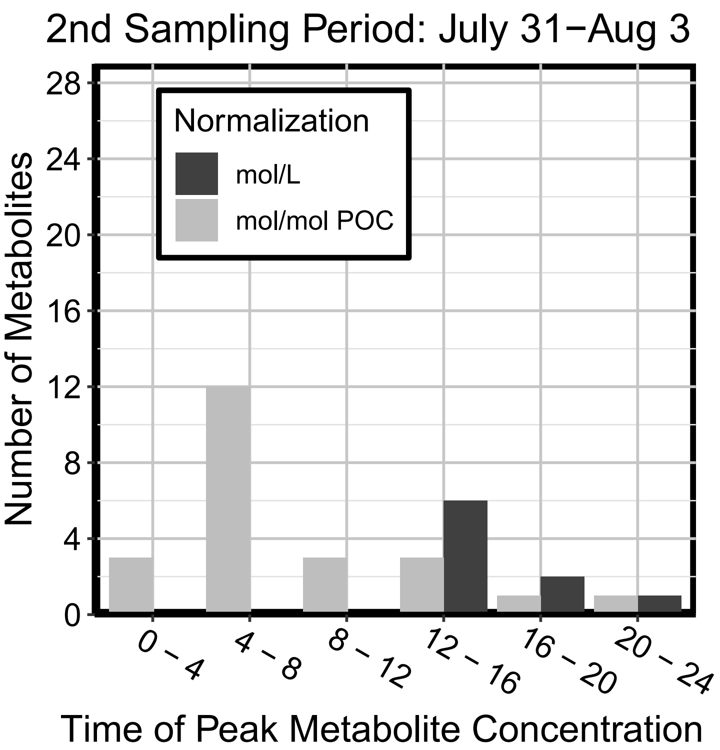


Supplemental Figure 6. Time of day that compounds peak in the second sampling period (July 31 - August).
