## Supplemental material for "Diel Oscillations of Particulate Metabolites Reflect Synchronized Microbial Activity in the North Pacific Subtropical Gyre"

### Supplemental Methods

#### *Metabolite sample extraction*

Metabolites were extracted as in as in Boysen et al.(1). Briefly, frozen filters were cut into small pieces and put into bead beating tubes with silica beads, heavy isotope-labeled internal standards (Table S1), and cold aqueous (50:50 methanol:water) and organic solvents (dichloromethane). The samples were shaken on a FastPrep-24 Homogenizer for 30 seconds and chilled in a -20 ˚C freezer, repeated for three cycles. The organic and aqueous layers were separated by spinning samples in a microcentrifuge at 5,000 rpm for 90 seconds at 4 ˚C. The aqueous layer was removed to a new combusted glass centrifuge tube. The remaining organic fraction was rinsed three times with 50:50 methanol:water. All aqueous rinses were combined with the original aqueous extract and dried down under $N_2$ gas. The remaining organic layer was transferred into a clean glass centrifuge tube and the bead beating tube was rinsed two more times with cold organic solvent. The combined organic rinses were centrifuged, transferred to a new tube, and dried under $N_2$ gas. Dried aqueous and organic fractions were re-dissolved in 400 $\mu$L of water and 400 $\mu$L of 1:1 water:acetonitrile, respectively. Isotope-labeled injection standards were added to both fractions (Table S1).

*Metabolite data acquisition*

LC-MS parameters were as in Boysen et al.(1). Data for most compounds were collected on a Waters Xevo TQ-S triple quadrupole (TQS), but for compounds outside the linear range or which were originally detected in sub-optimal polarity, a Thermo QExactive HF (QE) mass spectrometer with ESI was used. Chromatography and mass spectrometry methods follow those reported in Boysen et al.(1), and details are provided in the supplemental methods. The organic fraction was analyzed using reversed phase chromatography (Waters Acquity UPLC HSS Cyano column, 1.8 $\mu$m particle size, 2.1 mm x 5 mm) and the aqueous fraction was analyzed with both reversed phase and hydrophilic liquid interaction chromatography (HILIC - SeQuant ZIC-pHILIC column, 5 mm particle size, 2.1 mm x 150 mm, from Millipore). We monitored 101 compounds with reversed phase and 110 compounds with HILIC.

Data collected on a Thermo QExactive HF (QE) with ESI was used for nucleic acids and nucleosides (adenine, guanine, thymine, cytosine, adenosine, guanosine, thymidine, cytidine) and compounds which were overloaded on the triple quadrupole (glycine betaine, homarine, phenylalanine, DMSP) but which were within the linear range on the QE. On the QE, for HILIC, a full scan method with polarity switching was used with 60,000 resolution. For RP, positive ionization mode was used with a resolution of 120,000. For the QE data, proteowizard was used to convert .raw files to .mzxml(2).

Peak integrations were performed using Skyline for small molecules(3). Isoleucine and leucine did not always chromatographically separate and were treated as a single metabolite, (iso)leucine. Data were subjected to in-house quality control (QC) that removed misidentified compounds, removed compounds with a low signal to noise ratio (S/N < 4), and flagged compounds that were detected in the blanks (signal intensity in the sample must be greater than 3 times that in the blank). Compounds which were below the level of the blank in greater than 30% of samples (40 out of 132) were discarded. Compounds which did not pass the QC for 6 or more samples were discarded. For compounds which had fewer than 6 samples fail the QC, if the compound was detected in a sample but comparable to the blank, that data was retained. If the signal in a sample was less than the signal in the blank, a value equal to half the average blank value was filled back in to reflect the limit of detection.

*Metabolite data curation*

The normalization procedure for metabolomics data follows the Best-Matched Internal Standard method described in Boysen et al.(1) Pooled samples at full and half strength (diluted 1:1 with water, for the aqueous fraction, or solvent, for the organic fraction) were run after the samples were run in order to train the normalization algorithm. An improvement of 10% in the relative standard deviation (RSD) of the pooled sample was required in order to apply normalization. Compounds with a raw RSD of less than 0.01 were not normalized. The low cutoff was used to encourage normalization because the data were collected over a period of five weeks of instrument time; normalizing to an internal standard is an improvement over the instrument variability that we expect to influence mass spectral measurements over this time period.  All QE Orbitrap data were either used as raw data (no normalization) or, in the case of phenylalanine, normalized to its isotopologue internal standard, as the high resolution instrumentation experiences less ion suppression compared to that on the TQ-S instrument(1).

Prior to normalization, the peak areas of the internal standards in each sample were assessed to detect run quality and extraction efficiency. This assessment showed that a number of samples, mostly those collected as part of the second sampling period, clearly had half or twice the appropriate standard concentration added. (samples with internal standards adjusted in the $1^{st}$ sampling period: 5R1, 6R1, second sampling period: 26R2, 26R3, 27R1, 27R2, 27R3, 28R1, 28R3, 29R2, 30R1, 31R1, 32R3, 34R1, 34R3, 35R1, 35R2, 37R1, 39R1). Peak areas were adjusted according to these observations as well as laboratory notebook records for the samples which had incorrect concentrations. The periodicity analysis was done both with and without these replicates and the results are robust to this change. Additionally, samples collected between July $29^{th}$ 14:00 and July $30^{th}$ 02:00 did not have some of the standards added during sample processing. Data from the effected compounds were removed for those samples as normalization was not possible (as indicated in Table S2).

Euclidean distance of samples based on *z*-score standardized metabolite profiles showed that single replicates from two different samples (31R2, 41R1) in the second sampling period were outliers (> 3 standard deviations away from the mean average distance), so these samples were removed prior to further analysis.

*Quantification of select metabolites*

Metabolites with isotope-labeled authentic standards (see list in Table 3) were quantified using the following formula:

$$[M]_{SW} = \frac{PkA_M}{PkA_{IS}} * [IS]_{vial} * \frac{V_{vial}}{V_{filtered}}$$

where *M* indicates metabolite, *V* indicates volume, *IS* indicates internal standard, *SW* indicates seawater, and *PkA* is the integrated LC-MS peak area.

Isotope-labeled standards for trehalose and sucrose were purchased after the full sample set had been processed. A subset of samples (*n* = 19) was spiked with the internal standards and re-analyzed on the LC-TQS-MS, concentrations were determined as above, and a linear regression relating peak area to trehalose and sucrose concentration was fitted according to the formula:

$$[M]_{vial} = A \times (NormalizedPkA)$$

where *A* is a constant. This regression was used to calculate the concentrations of trehalose and sucrose in the rest of the samples that were not reanalyzed.

An authentic standard for 2,3-dihydroxypropane-1-sulfonate (DHPS) was obtained after the full sample set had been processed and the quantification of this compound using standard additions is described in Durham et al(4).

*Estimated Concentrations of Metabolites*

Approximate concentrations correcting for ionization efficiency and ion suppression were calculated using the equation:

$$Concentration = \frac{Area}{IE} * \frac{V_{vial}}{V_{filtered}} * \frac{1}{RF_{ratio}}$$

where *IE* is the ionization efficiency, calculated by taking the average peak area of a standard injected in water divided by the concentration of the standard. $RF_{ratio}$ is the response factor ratio published in Boysen and Heal et al. (2018), calculated by taking the ratio of the peak area of a standard injected in environmental matrix (less the ambient matrix signal, if applicable) to the peak area of the standard injected in water. Published values were used with the exceptions of taurine, chitobiose, and for nucleosides and nucleotides where we used values calculated from a different sample set also from near Station ALOHA, so with similar matrix effects.

We did not make estimates of concentration for five compounds: methylthioadenosine (MTA) in the aqueous fraction because we didn't have appropriate standards at the time of sample analysis, EPA, DHA, ergosterol, and trans retinal because insufficient isotope labeled internal standards were used for the lipid fraction.

*Particulate carbon and particulate nitrogen discrete sample analysis*

Samples for particulate nitrogen and particulate carbon were collected form the ship's underway flow through seawater, which sits at ~7 m water depth. Samples were collected onto combusted GFF filters. Filters were folded and stored in combusted aluminum foil at -80 ˚C until analysis, when they were thawed and dried overnight at 60 ˚C, balled into Ag and Sn boats, and analyzed using high-temperature combustion (1020 ˚C) on a ThermoQuest NC 2500 elemental analyzer.

*Eukaryotic metatranscriptome assembly*

Metatranscriptome assembly was conducted using Trinity(5) on the Pittsburgh Supercomputing Center's Bridges Large Memory system. Parameters include using in-silico normalization, a minimum k-mer coverage of 2, and a minimum contig length of 300. The raw assemblies were quality controlled with Transrate v1.0.3(6). To eliminate redundancy and duplication, the assemblies were merged and clustered at the 99% amino-acid identity threshold level with linclust in the MMseqs2 package(7).

*Eukaryotic metatranscriptome reference database*

We supplemented the curated MarineRefII reference database (http://roseobase.org/data/) with additional representatives of marine animal, fungal, protist and viral reference sequences (Table S5), totaling 641 marine eukaryotes and prokaryotes.

*Detecting periodicity and estimating time of peak concentration/abundance*

When conducting the RAIN analysis outliers removed during quality control were filled back in by taking an average of the other two replicates collected at that time point.

The time of peak concentration for each oscillating metabolite and peak abundance for each oscillating transcript was calculated by fitting a periodic oscillator according to the function:

$$[M] = A * \cos\left(\frac{2\pi}{24} * HourCollected\right) + B * \sin\left(\frac{2\pi}{24} * HourCollected\right)$$

as in Ottesen et al, 2014(8). The lag time between metabolites and transcripts coding for proteins that use or produce them was estimated simply by taking the difference between the two peak times, in hours.

*Multivariate analysis*

Metabolite concentrations were standardized to their *z*-score across the samples, having a mean of zero and standard deviation of one. Samples and metabolites were clustered based on Euclidean distances with the vegdist and metaMDS functions from the vegan R package (version 2.5-6)(9). Average-linkage clustering was calculated using the hclust function from the stats R package (version 3.6.1).

We used nonmetric multidimensional scaling (NMDS) to explore differences in samples based on their metabolite profiles. NMDS is an ordination technique that uses rank-order similarity between samples rather than absolute differences between samples to reduce dimensionality in the data(10). NMDS using a Euclidean distance matrix was chosen rather than principal components analysis (PCA) because NMDS relaxes the assumption that there are fewer variables than samples, which is not the case in metabolomics where data on the abundance of hundreds of compounds is collected for each sample. Additionally, NMDS avoids the assumption of linear relationships among variables.

NMDS was run with two ordination axes and 100 random starts. Significance of the stress value was tested with a Monte Carlo randomization test. Goodness of fit was assessed by correlating NMDS ordination results and Euclidean distances using both a non-metric and linear fit. Analysis of Group Similarities (ANOSIM) was used to test differences between the times of day that samples were collected as well as differences between the days of collection. ANOSIM was conducted on a Euclidean distance matrix with 1000 permutations. Pairwise ANOSIM was conducted to further clarify which times were significantly different.

*Phytoplankton culture growth conditions*

Cultures of *Crocosphaera watsonii* WH8501 were obtained from the Zehr lab at the University of California, Santa Cruz. They were grown with a square 12:12 light:dark cycle with 50 μmol photons $m^{-2}$ $s^{-1}$, at 26 ºC, and had an average growth rate of 0.17 $d^{-1}$. Cells were collected in mid-to-late exponential phase by gentle vacuum filtration onto 0.2 $\mu$m Omnipore filters using combusted borosilicate filter towers. Cells were enumerated using a Beckman coulter counter.

*Mixed layer depth*

Mixed layer depth was calculated from CTD profiles and is defined by a 0.03 kg/m3 density offset from 10 db.

*Supplemental calculation: Trehalose fueling nitrogen fixation*

To convert half a mol $N_2$ to one mol $NH_3$ requires 3 mol electrons and 8 mol ATP(11). One mol glucose can provide 24 mol electrons or 30–36 mol ATP, leading to a stoichiometry of between 2.08–2.35 moles $CH_2O$ to produce sufficient ATP and electrons to produce one mole of fixed $NH_3$(12). From Wilson and Aylward et al. *Crocosphaera* is responsible for 7.3 nmol N $l^{-1}$ $d^{-1}$ (13), which therefore requires between 15.2 and 17.2 nM of respired carbon. Our data has a range of 1.6 to 4.3 nM C in the form of trehalose drawn down every night, with a mean drawdown of 3.2 nM C in the form of trehalose. Using this range of required carbon and trehalose drawn down, we estimate that trehalose catabolism could fuel between 9% and 28% of $N_2$ fixation, with a mean value of 20%.

### Supplemental Results and Discussion

*Multivariate Analysis*

NMDS analysis of the samples produced a low stress value of 0.18 which was significant ($p < 0.01$), indicating that the sample scores are robust. The fit between the ordination distance and Euclidean distance had a non-metric $R^2$ value of 0.97 and a linear fit $R^2$ value of 0.88. NMDS analysis of the samples from the second sampling period (Figure S2) produced a low stress value of 0.17 which was significant ($p < 0.01$), indicating that the sample scores are robust. The fit between the ordination distance and Euclidean distance had a non-metric $R^2$ value of 0.97 and a linear fit $R^2$ value of 0.89. The overall similarity observed at 6:00 during the first collection period is not seen during the second collection period. NMDS and ANOSIM analysis of the samples from the second collection period and full dataset are unable to discriminate between samples collected at different times of day (Supplemental Figure 2), providing additional evidence that community synchrony, as illustrated by overall metabolite composition, weakened as sampling progressed.

*Metabolites lose diel periodicity in second sampling period*

Fewer metabolites had diel periodicity in the second sampling period (Figure S1), with 9 and 23 compounds exhibiting diel periodicity when analyzed as molar concentrations (nmol $L^{-1}$) and normalized to POC (nmol $\mu$mol $POC^{-1}$), respectively (Table S2, Figure S1). The reduction of diel oscillation in the second sampling period is seen in multivariate analyses as well, where in the second sampling period no sampling times clustered significantly (Figure S2).

Because these samples were processed over several weeks, it is possible that methodological issues arose and masked the diel oscillations during the second half of the sample set. We deem this unlikely because we still measure strong diel oscillations in some compounds over the course of the whole sample period. Overall there was not a drop in signal intensity from the instruments over the sample processing, and the increase in *Prochlorococcus* concentrations shows, if anything, that there would be more biomass rather than less biomass to produce a robust signal for the latter half of the data (Figure 1).
